# Deep-learning microscopy image reconstruction with quality control reveals second-scale rearrangements in RNA polymerase II clusters

**DOI:** 10.1101/2021.12.05.471272

**Authors:** Hamideh Hajiabadi, Irina Mamontova, Roshan Prizak, Agnieszka Pancholi, Anne Koziolek, Lennart Hilbert

## Abstract

Fluorescence microscopy, a central tool of biological research, is subject to inherent trade-offs in experiment design. For instance, image acquisition speed can only be increased in exchange for a lowered signal quality, or for an increased rate of photo-damage to the specimen. Computational denoising can recover some loss of signal, extending the trade-off margin for high-speed imaging. Recently proposed denoising on the basis of neural networks shows exceptional performance but raises concerns of errors typical of neural networks. Here, we present a work-flow that supports an empirically optimized reduction of exposure times, as well as per-image quality control to exclude images with reconstruction errors. We implement this work-flow on the basis of the denoising tool Noise2Void and assess the molecular state and three-dimensional shape of RNA Polymerase II (Pol II) clusters in live zebrafish embryos. Image acquisition speed could be tripled, achieving 2-second time resolution and 350-nanometer lateral image resolution. The obtained data reveal stereotyped events of approximately 10 seconds duration: initially, the molecular mark for initiated Pol II increases, then the mark for active Pol II increases, and finally Pol II clusters take on a stretched and unfolded shape. An independent analysis based on fixed sample images reproduces this sequence of events, and suggests that they are related to the transient association of genes with Pol II clusters. Our work-flow consists of procedures that can be implemented on commercial fluorescence microscopes without any hardware or software modification, and should therefore be transferable to many other applications.

## Introduction

Light microscopy is one of the most central tools of biological research, whether a biologist aims to get the first glimpse of a given cellular process or to quantitatively test the validity of hypotheses [1]. A specific area of application is the visualization of fluorescently labeled molecules. The design of such experiments is subject to inherent limitations [2, 3], requiring a trade-off between acquisition speed, signal-to-noise ratio (SNR), and prevention of photo-damage to the specimen [4]. These parameters cannot be optimized separately. For instance, to increase acquisition speed, exposure time must be reduced, leading to lower SNR [5, 6]. SNR can be recovered by, for example, increased power of the light used to excite fluorescence in the sample, resulting however in increased photo-damage.

While the experimental parameters during acquisition are subject to firm trade-off relationships, computational processing of images after acquisition can recover image quality. These approaches allow, for example, a further reduction of exposure times followed by computational reconstruction of low-SNR images. Conventional approaches for reconstruction of low-SNR images include projection methods [7], deconvolution filters [8, 9], and denoising methods [10, 11]. In the past decade, deep learning methods have become widely used in a variety of image processing applications, often outperforming conventional approaches [12]. In biological microscopy, deep learning has been successfully used for image classification [13–15], segmentation [16, 17], and restoration [18–21]. Initial deep learning approaches used standard deep networks to restore fluorescence microscopy images, requiring training data sets of matched low-quality and high-quality images. For example, networks can be trained on a reference data set with high SNR (“ground truth”), so as to restore matched images with low SNR (“noisy data”) [22]. One obstacle to the wide-spread application of such reconstruction approaches is the requirement for matched high-quality training data [23, 24]. These data are laborious or sometimes even impossible to obtain in a fashion that is sufficiently matched to noisy data. An alternative is provided by Noise2Noise (n2n) techniques, which enable the training of deep networks from matched pairs of noisy images [25, 26]. The requirement for any matched images is fully removed in the Noise2Void (n2v) technique, where learning and removal of noise are carried out based on a single noisy image data set [26, 27]. Reconstruction based on a single noisy data set also allows per-image training, thus compensating for day-to-day variability of, for example, fluorescence labeling or fine-adjustment of optical parts.

A second obstacle to the wide application of deep learning methods is the possibility of errors in the reconstructed fluorescence images [23, 24]. These errors manifest as deviations between the high-quality ground truth images and the images reconstructed from low SNR data. A dilemma arises, where the effective application of deep learning networks can only proceed without acquisition of ground truth data, but ground truth data are required to assure the experimenter that reconstruction is error-free. In this work, we develop a pragmatic work-flow for the quality-controlled adjustment and application of n2v for denoising in high-speed fluorescence microscopy. In this work-flow, for every acquired view of a given sample, a small data set with high-quality data is recorded to control reconstruction quality, followed by full time-lapse acquisition of only compromised data. We demonstrate the applicability of this work flow in the analysis of fluctuations in molecular clusters in live zebrafish embryos. Our analysis reveals a close coordination between post-translational modifications of RNA polymerase II (Pol II) and changes in the three-dimensional (3D) shape of these clusters on the scale of a few seconds. These observations are confirmed by an alternative experimental approach, where still images from chemically fixed cells are sorted based on an additional fluorescence marker for genes that transiently engage with the molecular clusters. Our approach provides a guideline for other microscopists interested in the quality-controlled application of ground-truth-free image reconstruction methods. The approach can be implemented on any fluorescence microscope typically used for time-lapse recordings without the need of software development or hardware control beyond the standard functionality.

## Results

### Quantification of image reliability and effective resolution in reconstructed microscopy images

The structural reliability and effective spatial resolution of reconstructed images can be assessed by a combination of widely used metrics. The structural reliability can be assessed via the structural similarity index metric (SSIM). SSIM quantifies the similarity between two images and returns a value between 0 and 1 [28, 29]. SSIM values close to 1 indicate that two images are very similar, lower SSIM values indicate images that are less similar. One application is the comparison of two images obtained with the same acquisition and post-processing steps, providing a quantification of the reliability of the obtained images. Using SSIM, we can, for example, demonstrate how changes in image acquisition settings, such as the reduction of exposure time, can compromise image reliability (Fig. S1). Applying n2v to pairs of reconstructed super-resolution microscopy images, we can illustrate how denoising can increase image reliability (Fig. 1A,B). SSIM can also be used to assess whether reconstructions of low-quality images obtained with, for example, low exposure times can approximate high-quality images (Fig. 1A,B). The assessment of image reliability via SSIM is, however, not sensitive to localized differences between images, as are typically introduced at edges during denoising procedures. Such local occurrences of unreliable reconstruction are readily detected by the local SSIM (Fig. S3) [30]. The combination of SSIM and local SSIM thus allows an assessment of image reliability based on paired images, as well as the similarity between a reconstructed and a corresponding high-quality image.

**Fig 1.**
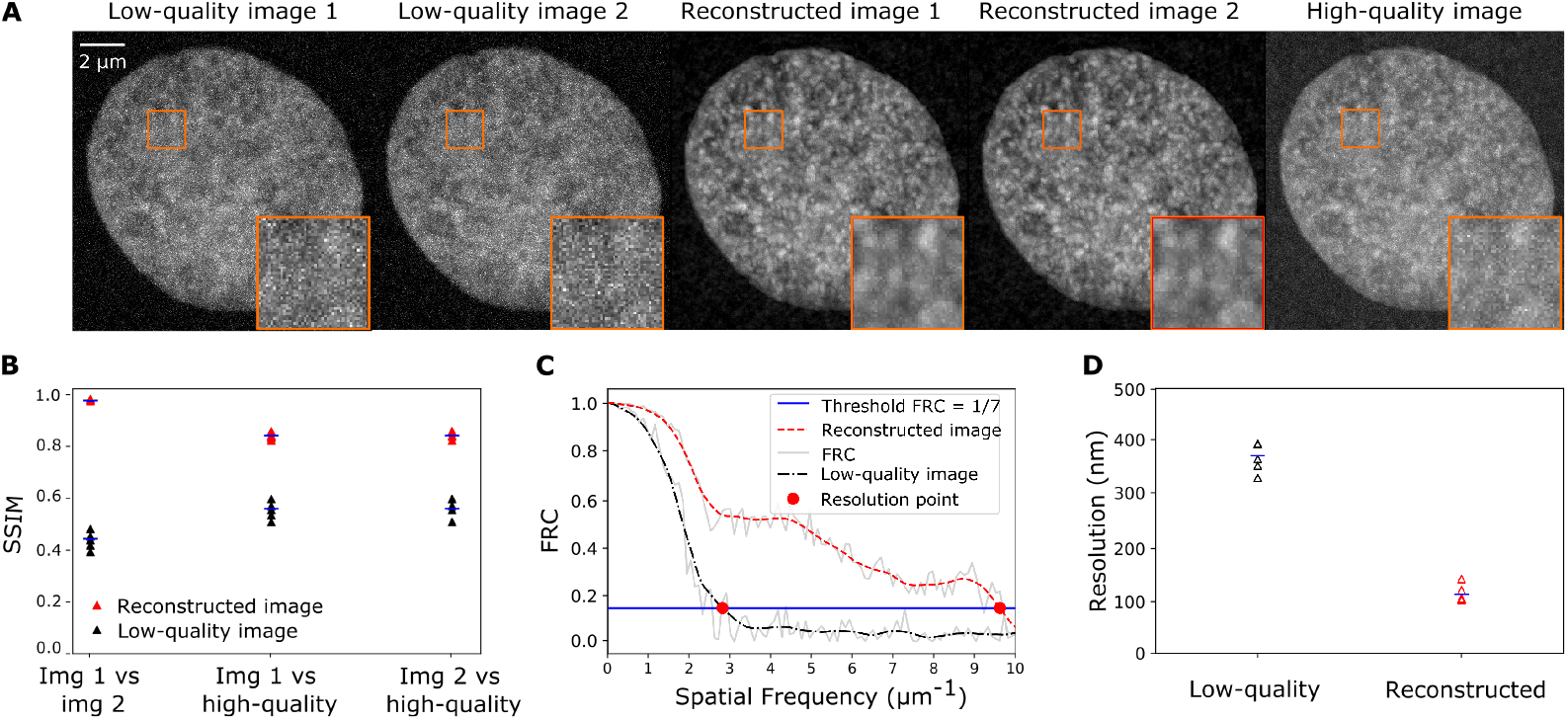
Metrics for the reliability and effective resolution in Noise2Void-reconstructed images. A) Representative micrographs of the DNA distribution in a nucleus in a fixed zebrafish embryo, recorded with a stimulated emission depletion (STED) super-resolution microscope. The same image plane was recorded twice at low quality, once at high quality, and two Noise2Void-reconstructed images were prepared from the low-quality images. B) SSIM values for pair-wise comparison (image 1 vs. image 2) and comparison against the high-quality image (image 1 vs. high-quality and image 2 vs. high-quality) for the low-quality images and the reconstructed images. C) FRC curves calculated based on a low-quality image pair and the corresponding reconstructed image pair. D) FRC-based effective resolution for four pairs of low-quality images and the corresponding pairs of reconstructed images.

A key aspect of performance in microscopy is the effective image resolution. The effective image resolution is determined by both the optical resolution of a given imaging instrument, and by the ratio of photons emitted by the structure of interest over polluting photons, often referred to as SNR. This effective resolution can be quantified via Fourier ring correlation (FRC) [31, 32]. FRC evaluates the similarity of a pair of images in frequency space, so as to determine the spatial frequency up to which the images are consistent with each other (Fig. 1C). The inverse of this spatial frequency is then taken as the effective spatial resolution (Fig. 1D). Applying the FRC metric to our super-resolution microscopy data reveals that, indeed, n2v-denoising can recover effective resolution in low-quality images (Fig.1D, Fig. S2). Taken together, SSIM and FRC can objectively assess image reliability and effective resolution in matched pairs of reconstructed images.

### Optimization of exposure time for high-speed time-lapse imaging

While denoising with n2v can, in principle, reconstruct images acquired with reduced exposure time (*t*_*exp*_), for a given experiment it is not known a priori just how far *t*_*exp*_ can be reduced while ensuring a sufficient image reconstruction. To demonstrate how SSIM and FRC can guide the choice of *t*_*exp*_, we carried out live sample microscopy of cells obtained from buccal smears (“human cheek cells”) for a range of exposure times, *t*_*exp*_ = 20, 40, 70, 100, 150 ms (Fig. 2A). For each *t*_*exp*_, a n2v-network was separately trained on a pair of images and the effective resolution for these reconstructed images was assessed (Fig. 2B). For *t*_*exp*_ = 70 ms, of 70 ms or higher, an effective resolution of ~ 200 nm was attained for the reconstructed images (Fig. 2C). This resolution was not further improved by longer exposure times, but could not attained for shorter exposure times (Fig. 2C). This FRC-based assessment suggest *t*_*exp*_ = 70 ms as an optimal exposure time. We controlled the structural reliability of the reconstructed images by local SSIM, finding reconstruction errors for *t*_*exp*_ = 20 ms (Fig. S3A-G). Considering both the FRC and local SSIM results, all *t*_*exp*_ ≥ 40 ms seem structurally reliable, while only *t*_*exp*_ ≥ 70 ms allows maximal effective image resolution after image reconstruction. In this setting, the experimenter can therefore choose between faster acquisition (*t*_*exp*_ = 40 ms) or higher effective resolution (*t*_*exp*_ = 70 ms), all while ensuring a high certainty of structural reliability.

**Fig 2.**
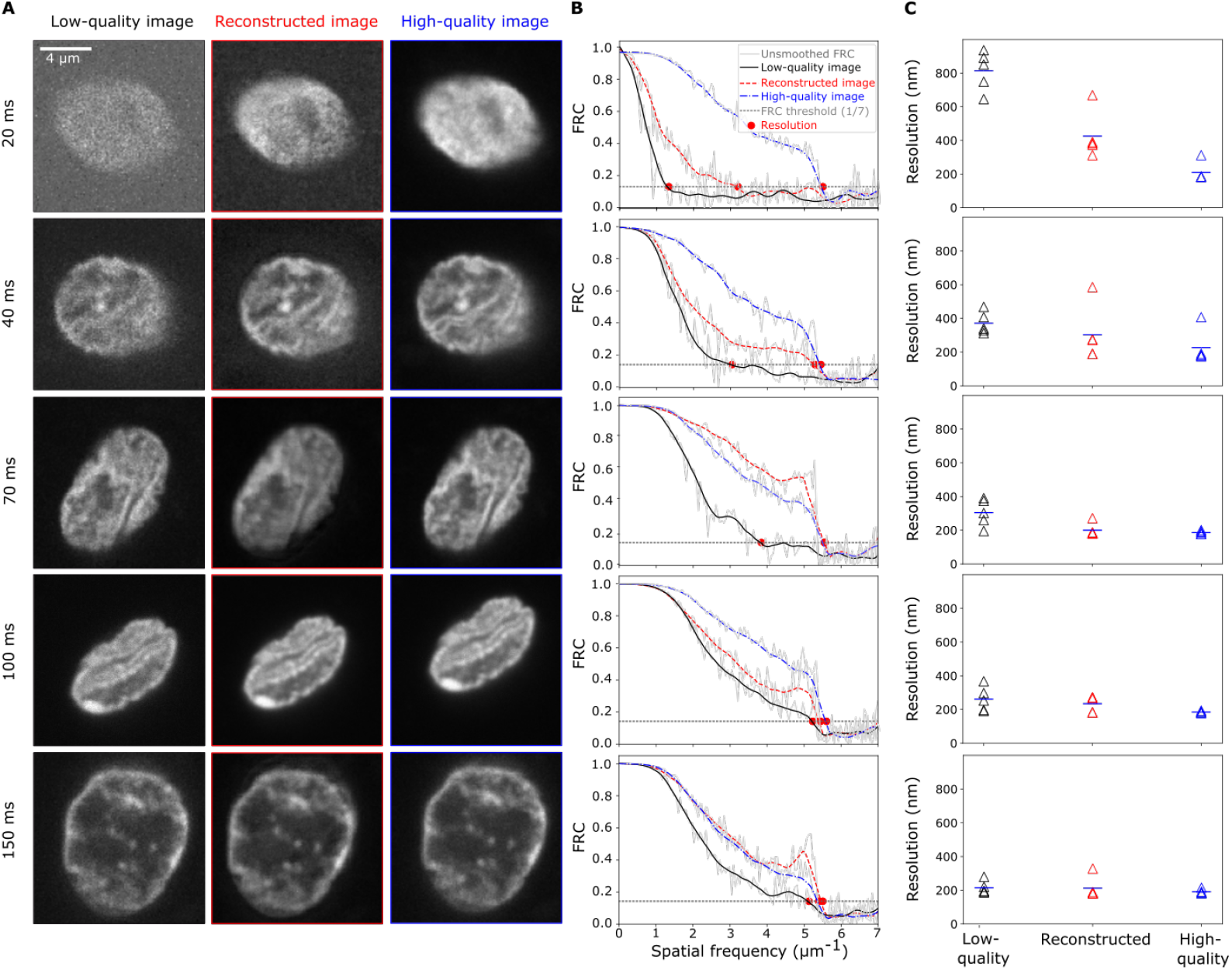
Metric-based estimation of how far image quality can be compromised to allow recovery of effective resolution by denoising. A) Representative micrographs of nuclei of human cheek cells for different camera exposure times (*t*_*exp*_, as indicated), all high-quality images were acquired at the same position but with an exposure time of 200 ms. Images are maximum-intensity projections, DNA was labelled by Hoechst 33342. B) FRC curves calculated from a pair of matched low-quality images, from a pair of reconstructed images, and a pair of high-quality images for the different *t*_*exp*_. C) Effective resolution for the indicated *t*_*exp*_, *n* = 5 nuclei per *t*_*exp*_, values are shown with mean.

### A two-phase acquisition protocol for quality-controlled denoising of time-lapse recordings

To integrate the metric-based assessment of n2v-processed images with the recording of high-speed time-lapse data, we propose an acquisition protocol that contains two distinct phases and is carried out at every position in a given sample (Fig. 3A). In the first phase (A, assessment), all image data required for the application of SSIM and FRC metrics are recorded (Fig. 3B). In particular, for each of the image planes that make up the acquired 3D volume, the following images are obtained: one low-quality image (*t*_*exp*_), two high-quality images recorded with a longer reference exposure time (*t*_*ref*_), followed by two more low-quality test images (*t*_*exp*_). In the second phase (B, time-lapse), a sequence of 3D volumes is acquired with only a single low-quality images for each of the image planes, reducing the time spent for the acquisition of a 3D volume (Fig. 3B).

**Fig 3.**
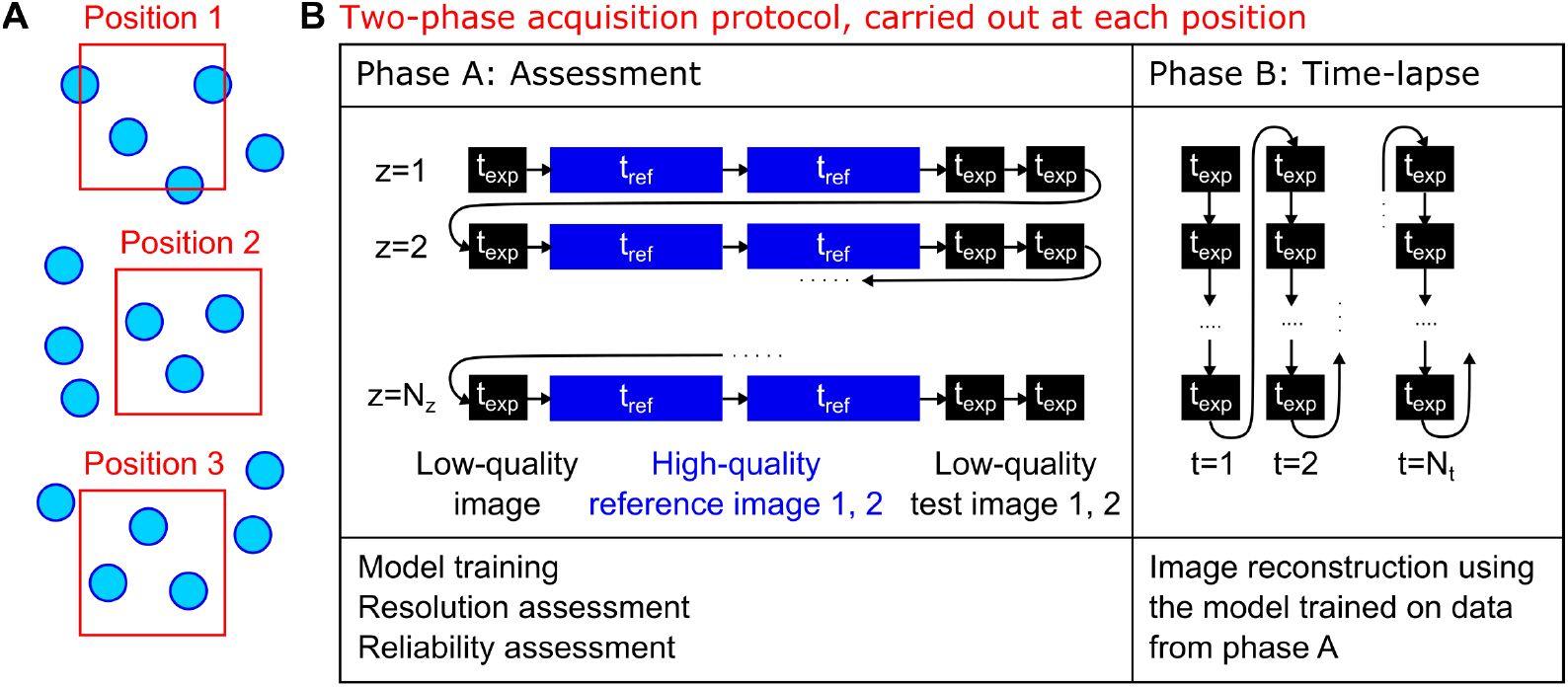
A two-phase acquisition protocol to combine acquisition of quality control images with high-speed time-lapse imaging. A) Image data were acquired at multiple positions in a sample, thus obtaining multiple viewpoints containing several objects of interest (nuclei, indicated in green). B) For each position, a sequence of two acquisition phases is carried out. In phase A, for each z position, a low-quality image, two high-quality reference images, and two low-quality test images are recorded. Low-quality images are recorded at a shortened exposure time (*t*_*exp*_), high-quality images at a reference exposure time resulting in images of the desired quality (*t*_*ref*_). Acquisition phase A obtains the images required for Noise2Void model training as well as the assessment of effective image resolution and reconstruction errors. In phase B, only single low-quality images are recorded with the shortened exposure time (*t*_*exp*_), resulting in an increased rate of acquisition compared to acquisition with full exposure time (*t*_*ref*_). Acquisition phase B obtains only low-quality images, which are reconstructed after the experiment is completed.

The data acquired by this two-phase acquisition protocol allow a comprehensive quality control assessment for every recorded position. Specifically, we first train a n2v network for each position, with which we reconstruct the low-quality test images 1 and 2. We can then assess the effective resolution using the FRC metric and additionally control the reconstructed image for reconstruction errors using SSIM and local SSIM. For positions where a sufficient effective resolution is achieved by the reconstruction, and a sufficiently low level of reconstruction error is found, the trained n2v-network is then applied to the time-lapse data from phase B, thus providing n2v-reconstructed time-lapses with per-position quality-control.

### High-speed imaging reveals coordinated changes of phosphorylation and morphology of RNA polymerase II clusters

To demonstrate the applicability of our proposed protocol for quality-controlled n2v-supported live imaging protocol, we attempted to visualize changes in the molecular state as well as the 3D shape of macro-molecular clusters enriched in Pol II. To this end, we recorded microscopy images from live zebrafish embryos with an instant-SIM microscope [33]. We visualized Pol II that is recruited to macromolecular clusters (Pol II Ser5P) or has transitioned towards production of RNA transcripts (Pol II Ser2P) by fluorescently labeled antibody fragments (Fabs). These Fabs provide good sensitivity in zebrafish embryos and do not perturb embryonic development in any obvious fashion [34–36]. To establish exposure times, we first adjusted imaging parameters so as to obtain images that reveal cluster shape in the Pol II Ser5P channel on the microscope’s live display without any processing. We chose this reference exposure time as *t*_*ref*_ = 200 ms, resulting in an overall time of 6 s that is required to obtain a full 3D image stack. Using *t*_*ref*_ = 200 ms, we recorded image data in line with the two-phase acquisition protocol, with the phase B spanning a total time of 2 min. Specifically, we recorded data for four different exposure times (*t*_*exp*_ = 10, 20, 50, 100 ms, Fig. S4A-C). For all *t*_*exp*_, we achieve an effective resolution of 400 nm (lateral) or better after n2v-based reconstruction, which compares favorably to an effective resolution of approximately 700 nm in the high-quality images (Fig. S4D). Analysis by local SSIM suggests that reconstructions for *t*_*exp*_ ≥ 20 ms offer a reliability similar to a comparison between two high-quality images, reconstructions of images obtained with *t*_*exp*_ = 10 ms are prone to reconstruction errors (Fig. S4E). Accordingly, we selected images acquired with *t*_*exp*_ = 20 ms (effective lateral resolution ~ 400 nm) and *t*_*exp*_ = 50 ms (effective lateral resolution ~ 350 nm) for further analysis, which provided full 3D image stacks at a time resolution of 1 s and 2 s, respectively.

As previously observed, clusters seen in the Pol II Ser5P channel were persistent during the entire phase B acquisition period [36]. The n2v-processed Pol II Ser5P time-lapse images were thus segmented to detect Pol II-enriched clusters, each cluster was then tracked over the whole time-lapse based on spatial proximity in consecutive time points (Fig. 4B). Based on the Pol II Ser5P-derived segmentation masks, Pol II Ser5P and Ser2P intensities as well as shape quantifiers (elongation and solidity) could be determined for each time point (Fig. 4A). The resulting time courses exhibit fluctuations, and the question arises whether a systematic relationship exists between the different quantities (Fig. 4C). Indeed, cross-correlation analysis that was anchored on cluster elongation reveals a systematic relationship (Fig. 4D). The cross-correlation analysis reveals an initial increase in Pol II Ser5P intensity, followed by a transient increase in Pol II Ser2P intensity ~ 5 s later, and a transient decrease in Pol II Ser5P intensity another ~ 5 s later. These changes are accompanied by an initial rounding up of clusters (solidity increase), followed by transient unfolding (solidity decrease) ~ 10 s later. These cross-correlation analysis results are obtained at both *t*_*exp*_ = 50 ms (Fig. 4) and *t*_*exp*_ = 20 ms (Fig. S5), indicating that our findings are not mere coincidence. Our observations are representative of a stereotypical sequence of events, which occurs repeatedly and is therefore detected by the cross-correlation analysis: Pol II Ser5P intensity increases and the cluster rounds up due to the rapid recruitment of Pol II to a given cluster, Pol II Ser5P intensity decreases and Pol II Ser2P intensity increases as some of the recruited Pol II proceeds into transcript production, while the cluster gets elongated and unfolded due to a shape perturbation resulting from transcript production. Previous work indicates that transcribing Pol II and the resulting nascent RNA transcripts induce distinct rearrangements in molecular clusters, explaining a potential source for the shape perturbation [35–38]. Notably, these works suggest that changes in Pol II state and cluster organization come about due to transient engagement of genes with Pol II-enriched clusters, leading to the induction of genes and to their release from these clusters.

**Fig 4.**
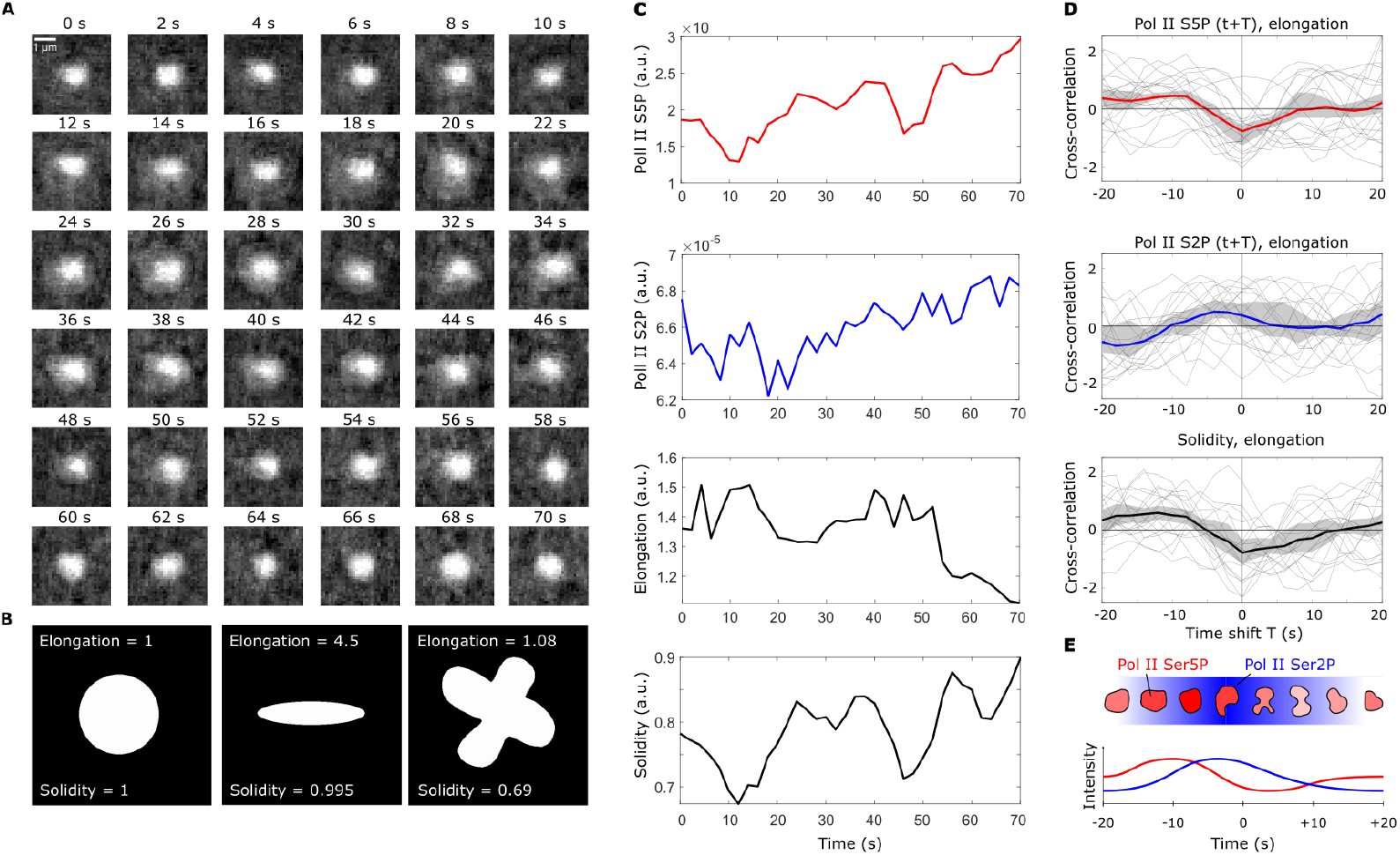
Noise2Void-accelerated imaging reveals coordinated changes in shape and phosphorylation levels of RNA polymerase II clusters on the scale of seconds. A) Representative series of time-lapse images showing a single RNA polymerase II cluster in the Pol II Ser5P channel (single image plane from the middle z position of the cluster, exposure time *t*_*exp*_ = 50 ms, effective time resolution for full 3D volume acquisition of 2 s) The Pol II Ser2P channel is not shown because only average intensity, not shape was quantified from this channel. B) Example shapes to illustrate how elongation and solidity represent object shape. C) Time courses of Pol II Ser5P intensity, Pol II Ser2P intensity, elongation, and solidity for the example time-lapse shown in panel A. D) Cross-correlation analysis of the temporal coordination of Pol II Ser5P intensity, Pol II Ser2P intensity, and solidity with elongation. Gray lines indicate the time-shifted correlation for single cluster time courses, thick lines indicate the mean, and the gray region the 95% bootstrap confidence interval. Analysis based on *n* = 30 tracked clusters, recorded from one sphere stage embryo. E) Summary of the coordinated changes in phosphorylation and cluster shape suggested by the cross-correlation analysis. A stereotypical sequence of events can be seen: cluster Pol II Ser5P intensity transiently increases (red) and the cluster becomes rounder, then cluster Pol II Ser2P transiently intensity increases (blue), until finally the cluster transiently unfolds and becomes elongated.

### Pseudo-time analysis from fixed sample images also detects coordinated changes in phosphorylation and cluster shape

To verify the conclusions obtained by the fluctuation analysis, we assessed changes in cluster state by an independent approach based on the interaction with a gene. Specifically, we fixed zebrafish embryos in the sphere stage, and fluorescently labeled a panel of 8 genes as well as Pol II Ser5P and Pol II Ser2P (Fig. 5A). Fixation of samples prevents live imaging, thus removing the temporal information from the images. In exchange, images with distinctly higher signal can be obtained without the need of n2v-processing, and the location of the labeled gene can be used as additional information that is not available in our live imaging data. The analysis of the obtained image data was therefore based on gene-Pol II cluster interaction pairs. An interaction pair is constructed by the detection of the location of a labeled gene, and by logical association of this gene with the Pol II Ser5P cluster that is closest in space (Fig. S6A,B). For each interaction pair, fluorescence intensities of the gene, fluorescence intensities of the the Pol II cluster, distance between both objects, and shape properties of the Pol II cluster were combined into a vector representing the interaction pair. Principal component analysis of these pairs revealed a cyclical pattern, based on which a pseudo-time coordinate was constructed (Fig. 5B and Fig. S6C). The assignment of a pseudo-temporal order to image data obtained from fixed samples has been used previously, for example for the nanoscale assessment of endocytosis [39, 40]. Ordering the interaction pairs along the pseudo-time coordinate allowed the extraction of time-shifted correlations (Fig. 5C), which directly mirrored those we obtained from on our live imaging data (Fig. 4D). We suspected that the location of a gene that interacts with Pol II Ser5P clusters provides the crucial information for successful pseudo-time reconstruction (Fig. S7, genes *foxd5, klf2b, zgc:64022*). Indeed, when we attempted pseudo-time reconstruction on the full panel consisting of eight genes, we found that for genes that only rarely come close to Pol II Ser5P clusters, the pseudo-time approach failed to reproduce the correlation analysis results (Fig. S7, genes *vamp2, ripply1, drll*.*2, gadd45ga, iscub*). In the case of successful pseudo-time reconstruction, our results suggests that a gene visits a Pol II Ser5P cluster in close coordination with changes that occur in the Pol II cluster. Specifically, genes engage in close contact when cluster Pol II Ser5P intensity increases, and detach at a time when clusters undergo transient elongation (Fig. 5A,B and Fig. S7, genes *foxd5, klf2b, zgc:64022*). The time-scales of this interaction can be estimated by a comparison of the distance between the cross-correlation maximum and minimum in the cluster Pol II Ser5P signal (~ 50 steps in pseudo-time, corresponding to ~ 10 s in the cross-correlation analysis based on live-imaging results) and the total number of observed interaction pairs (169, 186, 191 for *foxd5, klf2b, zgc:64022*, respectively), implying in an average duration of ~ 36 s between two consecutive interaction events. To conclude, the correlation analysis based on pseudo-time reconstruction provides an independent confirmation of the coordination between Pol II phosphorylation levels and cluster shape obtained by n2v-supported live imaging. This agreement suggests that these two approaches provide complementary views of the same, stereotyped sequence of changes in molecular properties and the shape of Pol II clusters.

**Fig 5.**
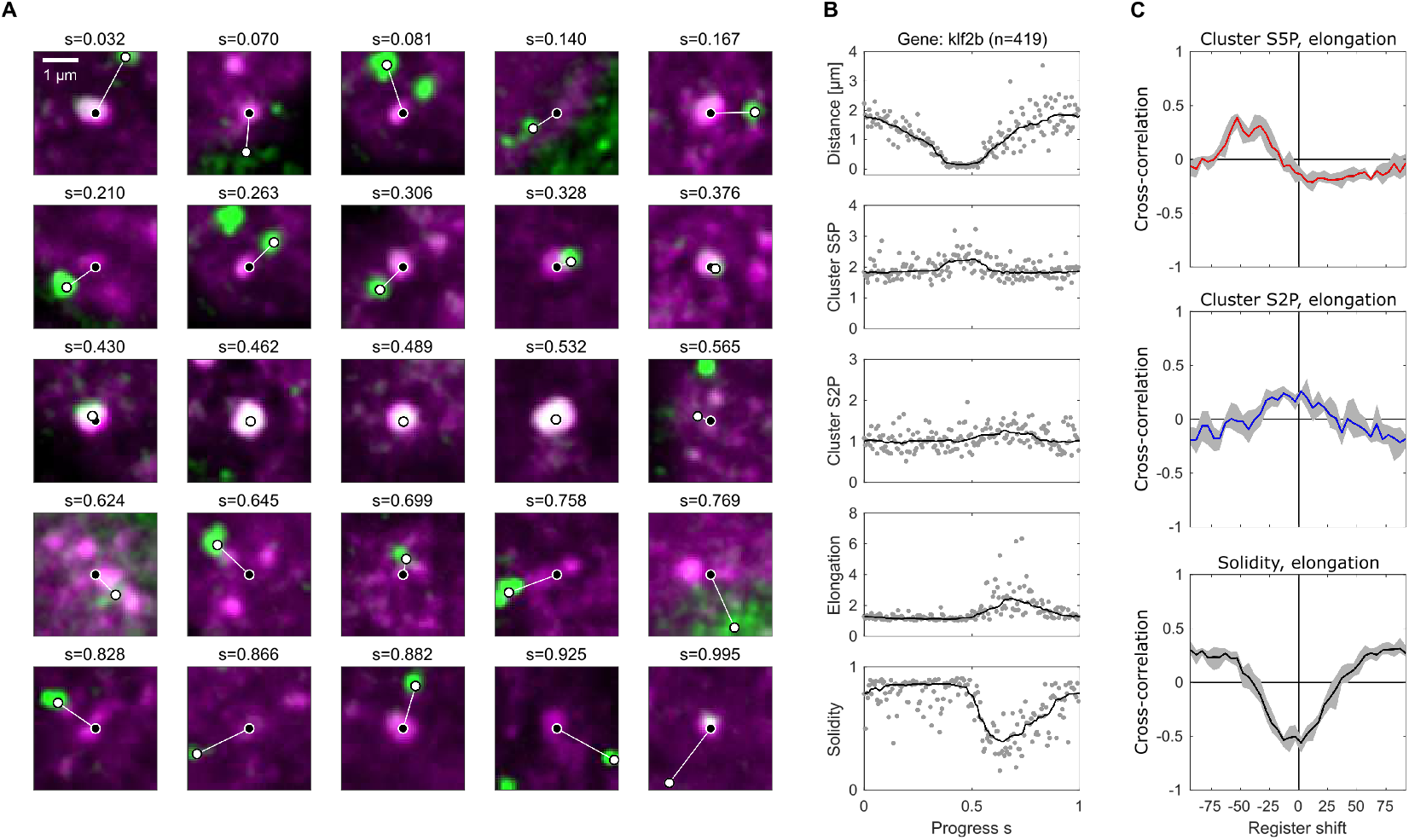
Pseudo-time analysis of data from fixed embryos relates transient engagement and activation of a gene to the phosphorylation and shape changes observed in live embryos. A) Example images of Pol II Ser5P (magenta signal) clusters sorted by a pseudo-time progress coordinate (*s*, periodic, defined on the interval [0, 1)), which is calculated based on interaction with the gene *klf2b* (green represents oligopaint fluorescence *in situ* hybridization signal for *klf2b*). Center positions (weighted centroid) are indicated for the Pol II Ser5P cluster (white circle with black filling) and the gene (black circle with white filling) and connected with a white line for illustration. For details of the reconstruction, see Fig. S6. For an overview containing all eight genes that were assessed, see Fig. S7. B) Pol II Ser5P and Ser2P intensity, elongation, and solidity of Pol II Ser5P clusters sorted by pseudo-time *s*. A total of *n* = 186 clusters from *N* = 4 independent samples, obtained in two independent experiments, were included in the analysis. C) Cross-correlation analyses for different register shifts in the coordinate *s*, the register shift is in units of data points by which the coordinate *s* was shifted. Gray regions indicate 95% bootstrap confidence interval.

## Discussion

In this study, we describe how the quality of images that are reconstructed by deep-learning algorithms can be controlled for, addressing the specific case of unsupervised denoising by Noise2Void. We implemented our approach of quality control towards the acceleration of high-speed imaging, where camera exposure times are reduced and the resulting loss of signal quality is recovered by n2v-denoising. We then apply our approach to the example of imaging the molecular state and the shape of RNA polymerase II clusters in live zebrafish embryos. Our work illustrates how, in a practical application setting, the performance improvements from deep-learning algorithms in fluorescence microscopy can be combined with a high level of confidence in the reconstructed images.

We specifically apply our quality-control approach to an unsupervised denoising technique, the deep learning-based tool Noise2Void [27]. Currently, reconstructions that map from noisy to high-quality data on the basis of paired training image data offer the highest reconstruction performance [22]. In many practical settings, such pairs of noisy and high-quality images cannot be obtained. An alternative is offered by reconstructions based on matched pairs composed of noisy images only (Noise2Noise [26]). Further developments now offer the possibility to reconstruct high-quality images directly from single noisy images (Noise2Self [41] and Noise2Void [27]). A self-supervised approach seems ideally suited to reconstruction tasks where fluorescence labeling exhibits strong variability, optical components are changed between different experiments, or sample properties vary on a day-to-day basis. These characteristics are typical of biological microscopy applications, highlighting the applicability of self-supervised reconstruction methods in this area. A crucial assumption of self-supervised denoising approaches is that the noise in each pixel is an uncorrelated sample from the same probability distribution. Newer variants of these algorithms explicitly adjust the probability distribution of the noise to different parts of the image, thus improving the results where additional information on the noise characteristics is available [19, 42, 43]. Yet other variants model the structure of the signal itself [44]. These newer variants of self-supervised denoising could provide further improvements in reconstruction performance, while retaining most of the pragmatic applicability of self-supervised reconstruction methods.

We base our assessment of image quality on two metrics, (local) SSIM and FRC. More generally, metrics for image quality assessment belong to three main groups of functionality. The first group includes methods assessing the quality of images against a corresponding reference image (high-quality image). These methods are called full-reference, emphasizing the need for high-quality reference data [28, 45]. SSIM and consecutive similarity (CSS) metric, which is a variation of SSIM [46], are in this category. We used (local) SSIM, which provides an error map by structurally comparing the reconstructed image with the reference image, and based on that error map controlled for reconstruction defects. The second group, called reduced-reference, contains methods which are not using matched reference images, but rather general knowledge of properties and statistics that are typical of a set of reference images [47]. Natural scene statistics (NSS) is one major method in this category [45]. The underlying hypothesis of all NSS-based method is that all the original images are “natural” and that a distortion process introduces some unnaturalness that can be quantified by deviation from models of natural signals. Due to the day-to-day variability of the signals produced by fluorescence microscopy of biological samples, modeling natural signals appears challenging. The third category of image quality assessment methods is called no-reference, because quality assessment proceeds without a matched reference image or other prior knowledge [48]. Fourier ring correlation (FRC) is in this category and we used it to assess the spatial resolution of the reconstructed images. Based on the achieved spatial resolution, we could decide how far exposure times could be reduced while still supporting successful denoising. One tool that implements several of these metrics for the assessment of local anomalies in super-resolution microscopy data is SQUIRREL [49]. The quality scores and error mapping provided by SQUIRREL can, in principle, also be applied to images reconstructed by deep-learning methods.

The image acquisition protocol we propose consists of a phase during which all necessary data for quality control are collected for a single time point (phase A), followed by high-speed time-lapse imaging with compromised image quality (phase B). This protocol seems appropriate for the acquisition of short bursts of images, where the main limitation lies in how many images can be acquired in a short amount of time. For other imaging challenges, different protocols could be developed. In a different situation where, for example, photo-bleaching limits the acquisition of long time courses, excitation light levels could be reduced, and the compromised signal could be recovered by denoising. In such an experiment, quality control points could be placed at regular intervals over the course of acquisition. In a setting where, for example, sample structure or the level of fluorescence labeling changes significantly over the course of recording, a quality control phase at the beginning and at the end of the experiment might be advisable. Besides the implementation of additional control points in the experimental procedure, such extensions of our simple two-phase protocol would need no further modification to the quality control approach we used in our work.

Our live-sample microscopy recordings reveal a stereotypical sequence of events, where the Pol II recruitment and pause-release steps of transcriptional induction are closely coordinated with changes in the shape of Pol II clusters. While previous studies achieve high spatial or temporal resolution, our approach combines high resolution in time as well as in space. Our temporal resolution of 1-2 seconds for a full 3D stack is comparable to previous assessments of Pol II localization [50, 51]. These studies, however, do not monitor the specific phosphorylation states associated with Pol II regulation. Imaging of these phosphorylation states was previously performed with an effective time resolution of 1 min for a single gene [38] or 10 s for an engineered gene array [52] for the acquisition of full 3D volumes. By fitting of kinetic models of Pol II regulation, these studies suggest rates of promoter escape of 2-2.5 min and production of the first 1 kb of transcript length within 2.5 min (assuming an elongation rate of 0.4 kb per minute). Photobleaching experiments assessing endogenous Pol II combined with computational modeling indicated 2.3 s for initiation and 42 s of pausing at the promoter, as well as an elongation rate of 2 kb per minute [53]. Lastly, another study suggests that 6.3 s are sufficient for Pol II to loosely associate with an induced gene as well as proceed into elongation [50]. While these estimates for the duration of induction and pause-release imply a broad spectrum of kinetics, our estimates of 2-3 s for promoter escape and approximately 36 s for the duration of one complete gene-cluster interaction cycle fall within the previously estimated range for promoter escape and RNA production. Besides temporal coordination, also relative distances have been assessed, for example between Pol II clusters and nascent mRNA [38] and between enhancers, Pol II, and the transcription start site [54, 55]. In these studies, nascent mRNA is displaced one hundred to several hundred nm relative to sites harboring transcriptional regulators and recruited Pol II. This displacement is in line with our observations that genes that undergo elongation are located outside of Pol II Ser5P clusters. In contrast to previous work, our approach reveals the full shape of the Pol II Ser5P clusters. Taken together, the kinetics of single-gene induction suggested by our live-sample experiments seem in line with previous work, and the spatial organization of clusters and interacting genes directly correlates with previous work assessing relative distances of different components of the transcriptional machinery.

Our pseudo-time reconstruction revealed that the changes in Pol II phosphorylation and cluster shapes are temporally coordinated with the visit of genes to the Pol II clusters. Previous work suggests that the Pol II clusters in early embryonic development form on regulatory chromatin regions, including super-enhancers [36, 56, 57]. Accordingly, our data seem to directly show single genes that undergo transcriptional activation during a visit to Pol II-enriched cluster that contain regulatory chromatin regions. Different models for such enhancer-promoter communication in transcriptional control were proposed [58–60]. The stereotypical sequence suggested by our data fits most closely to a condensate hit-and-run model, where genes transiently interact with enhancer-associated condensates for transcription initiation, and leave from the condensate in association with the onset of transcriptional elongation [60]. A condensate hit-and-run model can also explain earlier observations suggesting cyclic interactions, where genes repeatedly engage with and depart from Pol II-enriched clusters [61]. Such a model also could support the proximity-dependent activation of *Shh* by its enhancer *ZRS* [62, 63]. The activation of genes by enhancers was also found to not require direct contact, but can occur over a distance of 200 nm or more [64, 65]. These observations, together with evidence in support of the condensate hit-and-run model, allow speculations about a liquid-bridge model of enhancer-gene communication. In such a liquid-bridge model, genes transiently become embedded within an enhancer-associated condensate, allowing the transfer of transcriptional machinery, including Pol II, to the gene promoter [60, 66, 67]. While previous work indicates that the onset of RNA production at newly activated genes results in their exclusion from the enhancer-associated condensates [37, 68–71], the initial engagement with the enhancer-associated condensates is less well understood. Such an engagement would, however, be naturally explained by the formation of small condensates at promoters. Such condensates could emerge, for example, at CpG-rich regions that are placed directly upstream of promoter regions of many developmental genes and were found to contribute to gene-promoter contacts in three-dimensional space [72].

## Materials and Methods

### Live imaging of primary cell culture of human cheek cell

Cells were obtained by a buccal smear with a P1000 pipette tip. Short-term primary cell cultures were then created by adding the pipette tip to a micropipette and pipetting up and down several times in 2 ml of PBS (Dulbecco’s formulation) with 0.8 mM CaCl_2_ and 4 *μ*M Hoechst 33342. 500 *μ*l of this primary cell culture were transferred to one well of an 8-well ibidi *μ*-Slide (8-well glass bottom #1.5 selected D263 M Schott glass). The ibidi slide was sealed with parafilm to prevent evaporation and incubated for 1 h at room temperature to ensure flattening of the cell nuclei before microscopy. Participants provided free and informed consent. Procedures were reviewed and accepted by the Karlsruhe Institute of Technology ethics committee. Raw image data were stored in an anonymous fashion and are not for public release.

### Zebrafish husbandry

All zebrafish husbandry was performed in accordance with the EU directive 2010/63/EU and German animal protection standards (Tierschutzgesetz §11, Abs. 1, No. 1) and is under supervision of the government of Baden-Württemberg, Regierungspräsidium Karlsruhe, Germany (Aktenzeichen35-9185.64/BH KIT). Embryos used for the different experiments were obtained through spontaneous mating of adult zebrafish. Collected embryos were dechorionated with Pronase, washed 3 times with E3 embryo medium, once with 0.3x Danieau’s solution, and subsequently kept in agarose-coated Petri dishes or 6-well plates in 0.3x Danieau’s solution at 28.5^*o*^C.

### STED microscopy of DNA in fixed zebrafish embryos

Following protocols from our previous work [73], sphere-stage zebrafish embryos were fixed overnight in 0.3X Danieu’s solution supplemented with 2% formaldehyde and 0.2% Tween-20 at 4°C, permeabilized for 15 min using 0.5 % Triton X-100 in PBS, washed three times with PBS supplemented with 0.1% Tween-20, and mounted in TDE-media supplemented with 10x SPY-595 DNA fluorescence stain under selected #1.5 glass cover slips. STED microscopy was performed using a Leica TCS SP8 STED microscope (Leica Microsystems, Wetzlar, Germany) with a 775 nm depletion line and a motorized-correction 93x NA 1.30 glycerol objective (HC PL APO 93X/1.30 GLYC motCORR). 100% 3D-STED depletion was used.

### Live imaging of RNA Pol II CTD phosphorylation in zebrafish embryos

One-cell-stage embryos were dechorionated with pronase in 0.3x Danieau’s solution and covalently labeled fragments of antibodies (Fab) were micro-injected into the yolk. In each embryo, 1 nl of Fab mix (0.2 *μ*l 1 % Phenol Red, 1.5 *μ*l A488-labelled anti-Pol II Ser2P Fab, 3.3 *μ*l JF646-labelled anti-Pol II Ser5P Fab, Fab stock cencentration approximately 1 mg/ml) was injected. The embryos were mounted for microscopy in 0.7% low-melting agarose in 0.3x Danieau’s solution in ibidi 35 mm imaging dishes (#1.5 selected glass cover slips) at 512-cell stage of development, images were acquired at the sphere stage.

### Oligopaint FISH and immunofluorescence in fixed whole embryos

Embryos were fixed in the sphere stage of development (4% formaldehyde, 0.2% Tween-20 in 0.3x Danieau’s, fixation overnight at 4°C), permeabilized (0.5% Triton X-100 in PBS, 15 min), washed in PBS with 0.1 % Tween-20 (PBST) for 2 minutes, and treated with 0.1 N HCl for 5 min. A sequence wash steps followed: twice with 1 ml 2x saline sodium citrate buffer with 1% Tween-20 (2xSCCT), once with 2xSCCT+50 % formamide for 2 min at room temperature, and once with preheated 2xSCCT+50% formamide at 60^*o*^C for 20 minutes. Liquid was replaced with a hybridiziation mix: 50 *μ*l formamide, 25 *μ*l 4x hybridization buffer (40 % dextran sulfate, 8xSSC, 0.8 % Tween-20), 2 *μ*l 20 *μ*g/*μ*l RNase A, 10 *μ*M oligopaint probes labeled with Alexa 594, and ddH_2_O added to reach a total volume of 100 *μ*l. Denaturation at 90 ^*o*^C for 3 min was followed by overnight hybridiziation at 37°C. Hybridization was followed by the following wash steps: four times with preheated 2xSSCT at 60^*o*^C and twice with 2xSSCT at room temperature, 5 min incubation time for each step. Before proceeding with the immunofluorescence protocol, the samples were additionally washed three times with 1 ml PBST for 5 minutes. Samples were blocked with 4% BSA in PBST, 30 minutes at room temperature, followed by incubation of primary antibodies (mouse anti-Pol II Ser5P (4H8, 1:300) and rabbit anti-Pol II Ser2P (EPR18855, 1:300)) in 4% BSA-PBST overnight at 4°C. Samples were washed three times for 5 min with PBST, once with 4% BSA-PBST, and again incubated overnight, 4°C with secondary antibodies (goat anti-mouse conjugated with STAR RED (1:300) and goat anti-rabbit conjugated with Alaxa 488 (1:300)) in 4% BSA-PBST. Finally, samples were washed three times with 1 ml PBST and mounted in Vectashield H-1000 under #1.5 selected cover glass.

An overview of the oligopaint probe sets is shown in table 2. Full oligo sequences and scripts used in probe design are provided as a supplementary file. The raw image data and analysis scripts are provided in the form of Zenodo repositories, see Data Availability statement.

### Staining of a repetitive region to establish fluorescence in-situ hybridization

To test the FISH procedure without the need of a full oligopaint library, a single probe against a repetitive region was designed. The zebrafish genome was screened for long repetitive regions using BLAST. A region containing 100 repeats on chromosome 25 was selected (repeat sequence: 5’-CCGACGCATCTTCGTGCTGG CTTACATACTCCGCTGCACC AATGACTTGAATTGCAGCCT TGGGCGTATGCTGCTC). Probes were produced by the same protocol used for actual oligopaint production, using a primer Alexa Fluor 594-conjugation at the 5’ end (Thermofisher). Primer sequences are shown in Tab. 1).

**Table 1.**
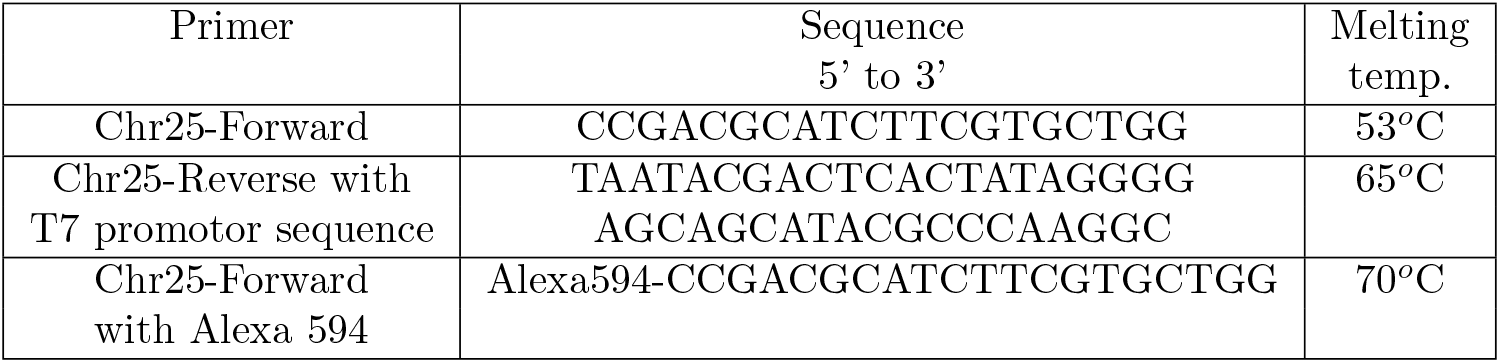
Primers used for PCR. Sequence (5’ to 3’) and melting temperature of each primer.

### OligoPaint probe library

Genes were chosen based on their Pol II Ser5P ChIP-Seq from Zhang et al., 2014 [74]. For each gene, Pol II Ser5P ChIP-Seq peaks in the promoter region (2 kb region upstream from the gene) was called using MACS2 [75]. For each peak, a p-value, q-value (Benjamini-Hochberg corrected p-value) and a signal value (fold enrichment of peak against background) was calculated. The peaks with p-value < 10^−5^, q-value < 10^−4^ and signal value > 3 were chosen. For these genes RNA-Seq data from White et al. 2017, were compared [76]. Only genes without high maternally provided RNA levels were considered, as indicated by low RNA counts for developmental stages preceding the high stage. Further, genes were hand-picked to cover a range of different RNA counts at the sphere stage. The scripts used for the gene selection process are provided as a supplementary file. OligoPaint libraries were designed using the OligoLego program [77]. Sequences of 32 nucleotides were mined from the zebrafish genome and are provided by the program (Tab. 2). Each probe set was designed to cover approximately 25 kb upstream and 25 kb downstream the gene, with density of 4 probes/kb where possible. The OligoPaint probe library was ordered from Twist Bioscience. The text file used to order the oligo pool is attached as a supplementary file. Associated amplification primers are listed in Tab. 3.

**Table 2.**
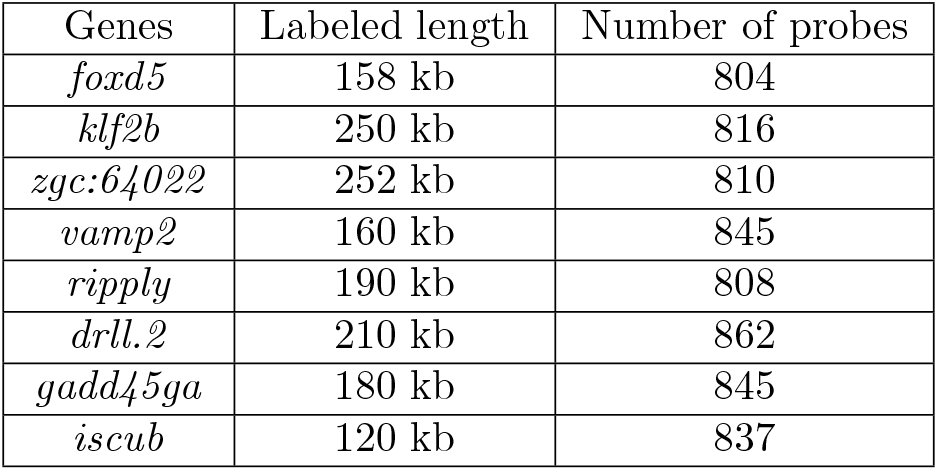
Genes chosen for OligoPaint DNA FISH, with the size of the labeled genomic region and number of OligoPaint probes covering the gene region.

**Table 3.**
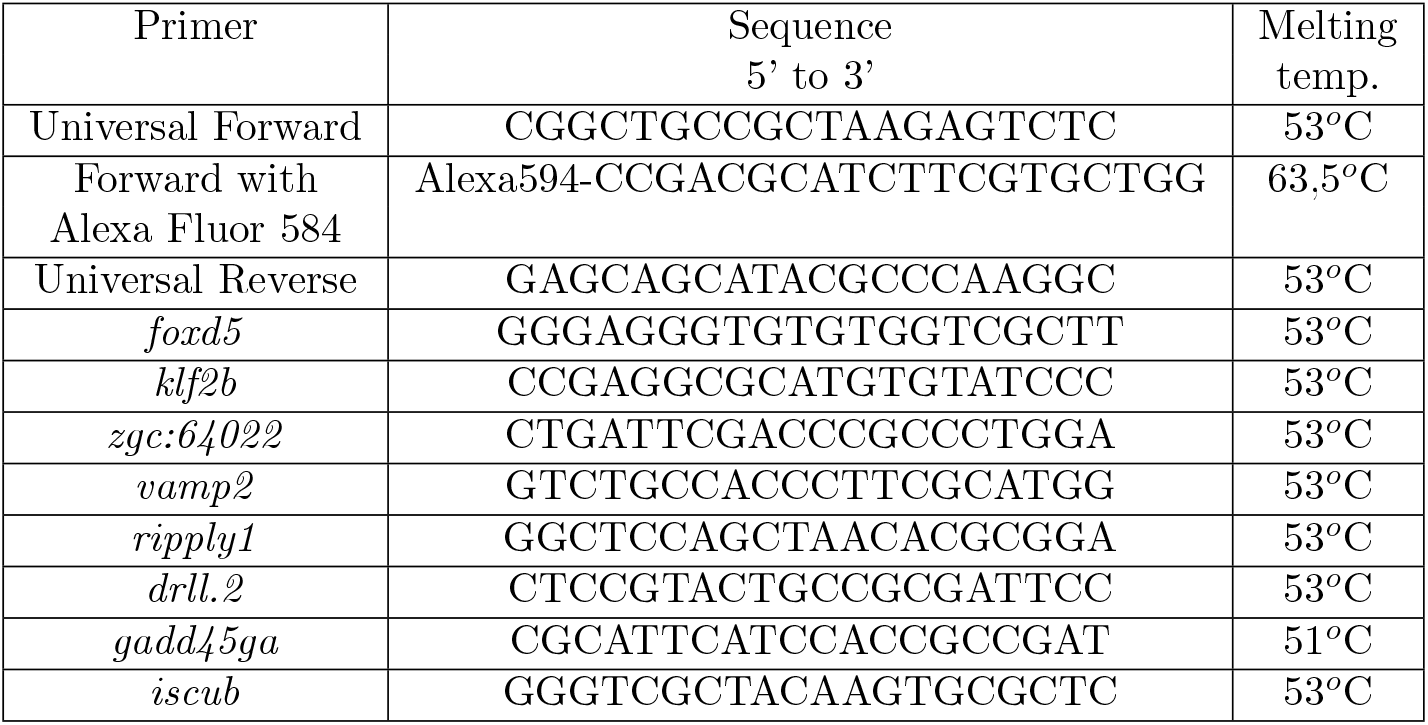
Sequence (5’ to 3’) of primers used for PCR for chosen genes for OligoPaint FISH and according melting temperatures.

### Instant Structured Illumination Microscopy (instant-SIM)

By using a VisiTech instant-SIM high-speed super-resolution confocal microscope, microscopy data were recorded from live human cheek cells as well as live and fixed zebrafish embryos. Microscopy data from live human cheek cells, live and fixed zebrafish embryos were recorded using a commercial implementation of the instant-SIM high-speed super-resolution confocal microscopy principle (VisiTech iSIM) [33]. The microscope was build on a Nikon Ti2-E stand. For live imaging a Nikon Silicone Immersion Objective (NA 1.35, CFI SR HP Plan Apochromat Lambda S 100XC Sil) and for fixed imaging a Nikon Oil Immersion Objective (NA 1.49, CFI SR HP Apo TIRF 100XAC Oil) were used. Laser at 405 nm, 488 nm, 561 nm and 642 nm were used for excitation. The acquisition settings were kept constant across all samples of a given experimental repeat.

#### Optimizing the exposure time for high-speed time-lapse imaging

In the process, on the one hand, time lapse recordings with a short exposure time of 150 ms, 100 ms, 70 ms, 40 ms or 20 ms were recorded every 5 seconds (21 loops), on the other hand, successive recordings with different exposure times (short, long, long, short, short) were recorded, in which “long” corresponds to an exposure time of 300 ms. For each exposure time a different cell nucleus was recorded. For each nucleus, on the one hand successive recording with different exposure times (short, long, long, short, short), on the other hand, time lapse images with the corresponding short exposure time for a total duration of 2 minutes, were recorded (see table 4). For each exposure time images from three embryos were recorded.

**Table 4.** Time-lapse parameters for high-speed imaging of Pol II clusters in live zebrafish embryos. total duration of time-lapse was 2 min.

| Condition | Exposure time | Full z-stack interval | Number of z-stacks |
| --- | --- | --- | --- |
| High-quality | 200 ms | 6 s | 21 |
| Shortened | 100 ms | 3 s | 41 |
| Shortened | 50 ms | 2 s | 61 |
| Shortened | 20 ms | 1 s | 121 |
| Shortened | 10 ms | 1 s | 121 |

### Noise2Void processing of STED microscopy data

For each nucleus, a pair of low-quality images a high-quality image were acquired. We first trained an n2v-network on the low-quality images 1 and reconstructed both low-quality images using this n2v-network.

### Noise2Void processing of cheek cell microscopy data

For each acquired position, we trained an n2v-network on the first low-quality image and reconstructed both low-quality images with the trained network. Procedures were reviewed and accepted by the Karlsruhe Institute of Technology ethics committee. Raw image data were stored in an anonymous fashion and are not for public release.

### Metrics for image assessment

The SSIM metric for the comparison of images *x* and *y* is given by

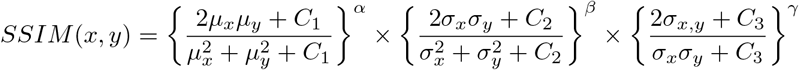

where *C*_1_, *C*_2_ and *C*_3_ are constants with the default value of 0.01 and 0.03, *C*_2_*/*2 respectively. With three constant *α, β* and *γ* (default values are (1, 1, 1)), the contribution of each term can be defined. *μ*_*x*_ and *μ*_*y*_ are the mean over each of the images’ pixel intensities, *σ*_*x*_ and *σ*_*y*_ are the standard deviations, and *σ*_*x,y*_ the covariance of the two images’ pixel intensity values. The first, second and the third term are respectively called luminance (mean), contrast (standard deviation) and structural (covariance). As the standard deviation of an image is not usually affected by denoising [46], in our experiment, we only focused on luminance and structural term and we investigated how luminance and structural terms of SSIM depend on the duration of photon collection in integrating versus averaging mode detectors. We observed that, for integrating detectors, the mean term influences the SSIM value, so that structural reliability cannot be directly compared if this term is included in the calculation of the SSIM value (Fig. S1A-D). For SSIM and local SSIM analysis, we therefore only consider the structural term, given by

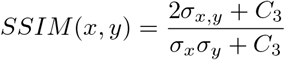

In local SSIM experiments, we used a Gaussian kernel with standard deviation of 12 pixels for weighting the neighborhood pixels around a pixel.

The Fourier ring correlation (FRC) analysis is based on the cross-correlation of two images in frequency space, and relies on the assumption that the two images are two independent reconstructions of the same object with independent noise realizations. The spatial frequency spectra of the two images are first divided into bins, which are in turn sorted by location within ring-shaped regions relative to the center of the Fourier spectrum polar coordinates. The FRC curve then is calculated based on the cross-correlation of the power values over all bins for a given ring radius, *r*, as follows:

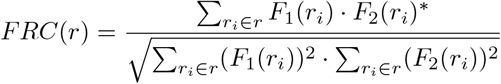

where *F*_1_, *F*_2_ are the Fourier transforms of two images and *r*_*i*_ refers to all frequency space bins that fall within a given ring radius *r*. The cut-off spatial frequency is the smallest *r* value for which the *FRC*(*r*) value drops below the widely used threshold value of 1*/*7. Up to the spatial frequency given by 1*/r*, the object is considered to be reliably resolved.

### Analysis of morphology fluctuations in RNA polymerase II clusters

Input images are recruited RNA polymerase II (Ser2P and Ser5P) in live zebrafish embryos visualized with antibody fragments (Fab) labelled with Janelia Fluor (Kimura Lab, Tokyo Tech), consisting of two channels, Pol II Ser2P and Pol II Ser5P, recorded by our proposed phase-AB imaging protocol (Fig. 3) with phase B spanning a time of 2 min. Data are specifically recorded for four different exposure times (*t*_*exp*_ = 10, 20, 50, 100 ms) (Fig. S4A).

#### Noise2Void denoising

We run the n2v script on Google Colab at the beginning of every exposure time for channel 2 (Pol II Ser5P) images. The patches which are given to the network are of size (16, 64, 64), while 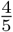 of the patches were used to train the network and 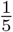 of all patches were used for the validation. We leave the network to be trained on each image for 70 epochs with the neighbourhood radius = 10. After assessing the n2v-processed images with FRC and local SSIM (Fig. S4), time-lapse Ser5P series recorded by 20 and 50 ms exposure time were selected and reconstructed using the trained network. The Pol II Ser2P time-lapse series left unchanged.

#### Median filter

After n2v-recovery of Pol II Ser5P series, to recover the signal a median filter of size 100 pixel is conducted on both Pol II Ser2P and Ser5P images.

#### 3D segmentation

We used an adaptive threshold based on the neighbourhood of size [101, 101] to segment each focus of Pol II ser5p images in 3 dimensions. Adapt threshold is a local threshold calculated based on the local mean intensity (first-order statistics) in the neighbourhood of each pixel. The Pol II Ser5P foci are segmented in 3 dimensions and the equivalent area in Pol II Ser2P images is also masked.

#### Foci tracking

Then, each focus is to be tracked over time. The distances between foci are used as the metric of tracking. Tracking was continued if a focus disappeared at *t*_*n*_ and again appeared at *t*_*n*+1_. We then extracted and saved the foci which lasted longer than 70 *s* for the further analysis.

#### 2D segmentation

We then picked the middle plane of each 3D focus and segment the middle plane in 2 dimensions. The plane which is used for 2D segmentation is rough of size (100 × 100) pixels. An adaptive threshold is again used for 2D segmentation (Fig. 4B and Fig. S5A).

#### Shape quantification

For each focus, we reported four properties (Ser2P mean intensity, Ser5P mean intensity, solidity, area and elongation) (Fig. 4C and Fig. S5B). If the focus lost at *t*_*n*_ and again appeared at *t*_*n*+1_, the average quantification of the focis at *t*_*n*−1_ and *t*_*n*+1_ is instead considered. The elongation property is calculated by the division of the major axis and the minor axis of the focus. For the calculation of Ser2p mean intensity property, the corresponding Ser5P segmented area in Ser2P images is considered. For Ser5P intensity property, the intensities of central pixels (rough of size 9 × 9 pixels) is averaged.

#### Correlation analysis

In this step, time lagged ([− 20, 20]) cross correlation between 3 pairs of (Ser5P intensity, elongation), (Ser2P intensity, elongation) and (Solidity, elongation) are quantified (Fig. 4D, Fig. S5C).

### Pseudo-time analysis from single time point fluorescence images

In this analysis, we work with still images of Pol II clusters, which were obtained from fixed zebrafish embryos. While these do not allow tracking Pol II clusters over time, the additional oligopaint fluorescence label can be used as information to sort images along hypothetical timeline. The pseudo-time analysis used for this sorting approach works under the assumption that, in cases where genes engage with Pol II clusters for transcriptional activation, a stereotyped sequence of events occurs that includes changes in Pol II phosphorylation, cluster shape changes, and association of the gene with a given cluster. Based on this assumption, the goal of this analysis is to reproduce such a stereotypical sequence by sorting Pol II cluster-gene pairs detected from an ensemble of still images

Nuclei were segmented by Otsu thresholding of blurred (Gaussian blurring, *σ* = 1.0 *μ*m) and background-subtracted (after Gaussian blurring, *σ*. = 10 *μ*m). Only nuclei with a volume greater than 40 *μ*m^3^ and a solidity greater than 0.7 were retained for further analysis. Pol II clusters were segmented inside each nucleus separately by robust background thresholding (2 standard deviations above intensity mean) after background subtraction (3.0 *μ*m) from the Pol II Ser5P channel. Only Pol II clusters with a volume greater than 0.03 *μ*m^3^ were retained for further analysis. Oligopaint-labeled genes were detected by robust background thresholding (6 standard deviations above intensity mean) of smoothed (Gaussian blurring, *σ* = 100 nm) and background-subtracted (Gaussian blurring, *σ* = 5 *μ*m) images of the oligpaint channel. Only objects with a volume greater than 0.05 *μ*m^3^ were retained for further analysis.

Gene-cluster interactions should be detectable by spatial proximity of a given gene to a Pol II cluster. The analysis thus connects any detected gene to the nearest neighboring Pol II cluster (Euclidean distance, Fig. S6B), resulting in a cluster-gene pair. These cluster-gene pairs are from here on treated as single observations, the remaining task is to sort these cluster-gene pairs into a coherent sequence. Only gene-cluster pairs with a cluster volume greater than 0.2 *μ*m^3^ were retained for further analysis.

The sorting of cluster-gene pairs is based on a mapping of correlated changes in the properties of cluster-gene pairs. Specifically, each cluster-gene pair is represented as a point an 8-dimensional feature space (ℛ^7^) defined by the gene and cluster properties (S6B). Application of a principal component analysis (PCA) to this ensemble of ℛ^8^ coordinates allows an effective reduction of dimensionality, and provides a mapping into distinct regions in the space spanned by the two first principal components (Fig. S6C). Plotting the projections of cluster volume and gene Ser5P level (check what the support lines actually are) into this PCA plot, an orthogonal coordinate system can be defined (Fig. S6C). Using only linear transformations (rotation and reflection), the cluster volume can be used as one axis of the coordinate system, and the gene Pol Ser5P placed to the left side of the plot. Inside this plot, data points can now be directly sorted in clock-wise order, providing a pseudo-time sorted dataset (Fig. S6C). This pseudo-time sorting is only successful when a gene engages in close contact with Pol II clusters with an increased frequency (centroid-centroid distance threshold for contact detection of 200 nm); genes that only engage Pol II clusters less often do not exhibit a useful pseudo-time sorting (Fig. S6D).

For the pseudo-time-sorted data points, a periodic progress coordinate *s* ∈ [0, 1) can be defined, assigning to each data point *i* a coordinate

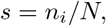

where *n*_*i*_ is the pseudo-time sorted index of the data point *i* in a data set consisting of a total of *N* data points. This progress coordinate can, in turn, be used to carry out cross-correlation analyses, based on a pseud-time shift Δ*s* (Fig. S7). For genes with high frequencies of engagement with Pol II clusters, this cross-correlation analysis closely reproduces the relationship between cluster morphology and Pol II Ser5 phosphorylation seen in our live imaging experiments (Fig. S7 *foxd5, klf2b, zgc::64002*). For genes with lower frequencies of engagement, the clear patterns of correlation between cluster elongation and Pol II Ser5 phosphorylation are not detected (Fig. S7 *vamp2, ripply1, drll*.*2, gadd45ga, iscub*).

## Supporting information

Scripts used for the selection of genes

Text file used to order the oligo pool

## Data Availability

Scripts and raw data are available at the following URLs.

Microscopy data and scripts for analysis of local SSIM and FRC in live zebrafish embryos: https://doi.org/10.5281/zenodo.5568871

Python scripts for assessment of different SSIM terms for integrating vs. averaging photon collection: https://doi.org/10.5281/zenodo.5569195

Scripts for SSIM and Fourier ring correlation analysis of Noise2Void-reconstructed STED images: https://doi.org/10.5281/zenodo.5569432

Microscopy data of RNA Pol II CTD phosphorylation in live zebrafish embryos: https://doi.org/10.5281/zenodo.5566880

Scripts used for morphology fluctuations analysis in Pol II clusters: https://doi.org/10.5281/zenodo.5569475

Matlab scripts used in the analysis of oligopaint-immunofluorescence image data: https://doi.org/10.5281/zenodo.5524939

Microscopy data for the gene *drll*.*2* : https://doi.org/10.5281/zenodo.5266592

Microscopy data for the gene *iscub*: https://doi.org/10.5281/zenodo.5266736

Microscopy data for the gene *vamp2* : https://doi.org/10.5281/zenodo.5266903

Microscopy data for the gene *gadd45ga*: ttps://doi.org/10.5281/zenodo.5268538

Microscopy data for the gene *foxd5* : https://doi.org/10.5281/zenodo.5266995

Microscopy data for the gene *klf2b*: https://doi.org/10.5281/zenodo.5268833

Microscopy data for the gene *ripply1* : https://doi.org/10.5281/zenodo.5268779

Microscopy data for the gene *zgc::64022* : https://doi.org/10.5281/zenodo.5268683

## Acknowledgement

This work is supported by the Helmholtz Association under the joint research school “HIDSS4Health – Helmholtz Information and Data Science School for Health” and the Helmholtz program Natural, Artificial, and Cognitive Information Processing (NACIP). HH was supported by HIDSS4Health, IM and LH were supported by the Priority Program Molecular Mechanisms of Functional Phase Separation of the German Science Foundation (DFG-SPP2191). Antibody fragments (Fab) for live imaging were kindly provided by the Kimura laboratory (Tokyo Institute of Technology). STED microscopy was conducted at the Karlsruhe Center for Optics and Photonics (KCOP). We thank Ralf Mikut for comments on our manuscript.

## Supplementary Materials

Scripts used for the selection of genes to be labeled by OligoPaint-FISH:

`Oligopaint_GeneSelection_ProbeDesign_Scripts.zip`

Text file used to order the oligo pool:

`OligopaintsToOrder_Oct2020.txt`

**Fig S1.**
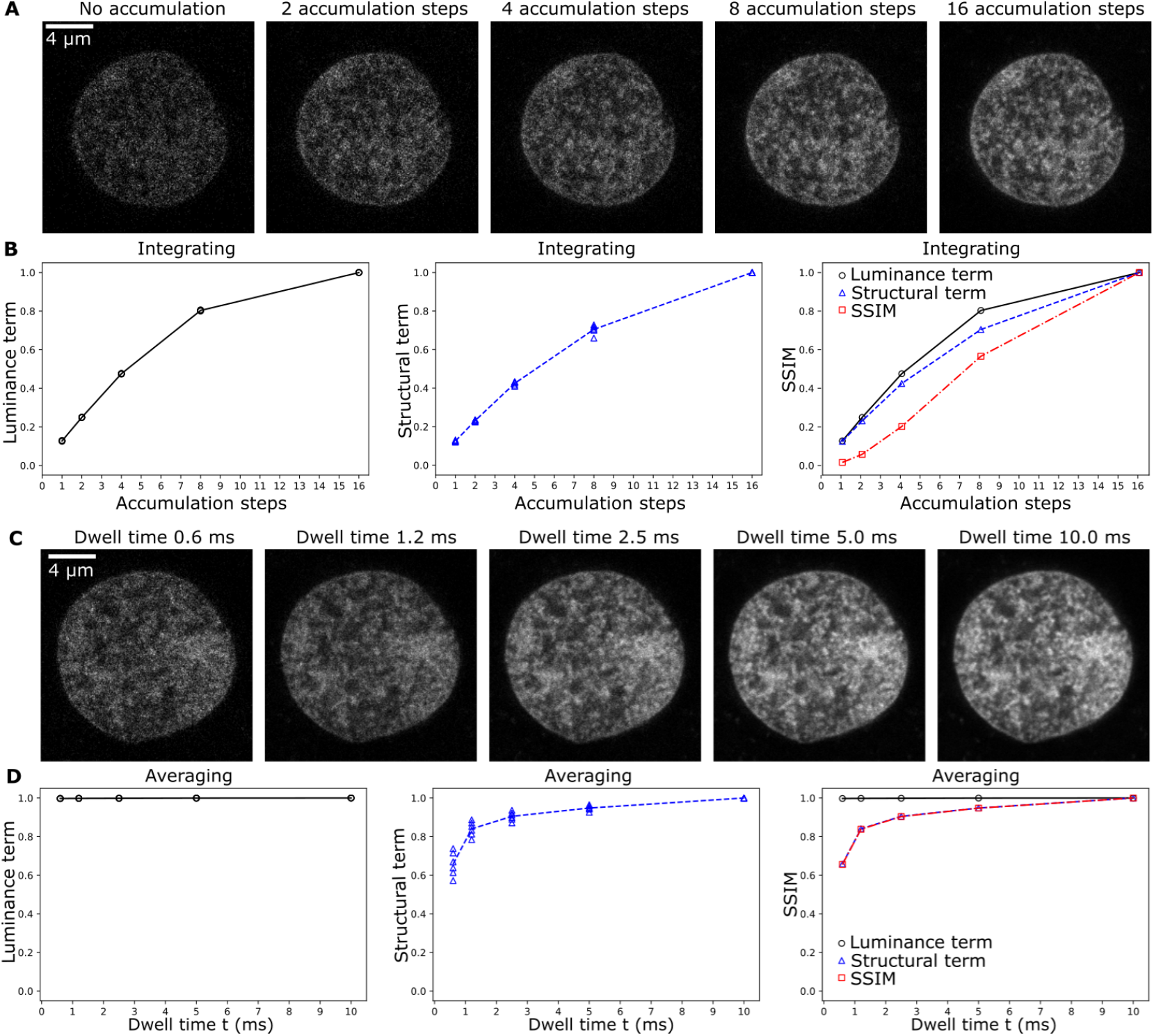
Structural similarity is a metric of proper structure. Figures are DNA distribution in a nucleus in a fixed zebrafish embryo, obtained by stimulated emission depletion (STED) microscopy. A) Depending on the microscope detector type and settings, photons can be accumulated over time without normalization (integrating detector) or with normalization (averaging detector). To illustrate how the different terms of the structural similarity index metric (SSIM) depend on the duration of photon collection in the integrating type detector, images were acquired with line-repeat scans in the accumulation mode, using increasing numbers of line repetitions (1, 2, 4, 8, 16), accumulation steps. B) In the integrating detector case, the luminance (mean) as well as the structural (covariance) term of the SSIM depend on the number of accumulation steps. Overall SSIM values contain contributions from both terms, so that an assessment of structural reliability would be obscured by changes in overall image intensity. SSIM values were calculated based on *n* = 6 images obtained by a reduced number of accumulation steps and a reference image obtained with the highest number of accumulation steps (16). Individual values are shown with the mean. C) To illustrate how the different terms of the SSIM metric depend on the duration of photon collection in the averaging detector type, DNA images were acquired with the detector is left open to collect photons at each pixel for longer times (dwell time *t*), then the photon count is normalized by the dwell time *t*. D) In the integrating detector case, the luminance term is constant and close to the value of 1.0. Only the structural term changes with increasing *t*, so that also the overall SSIM values directly reflect structural reliability for a given *t*. SSIM values were calculated based on *n* = 6 images obtained for a given *t*, combined with a matching image recorded with the highest *t* = 10 ms. Individual values are shown with the mean.

**Fig S2.**
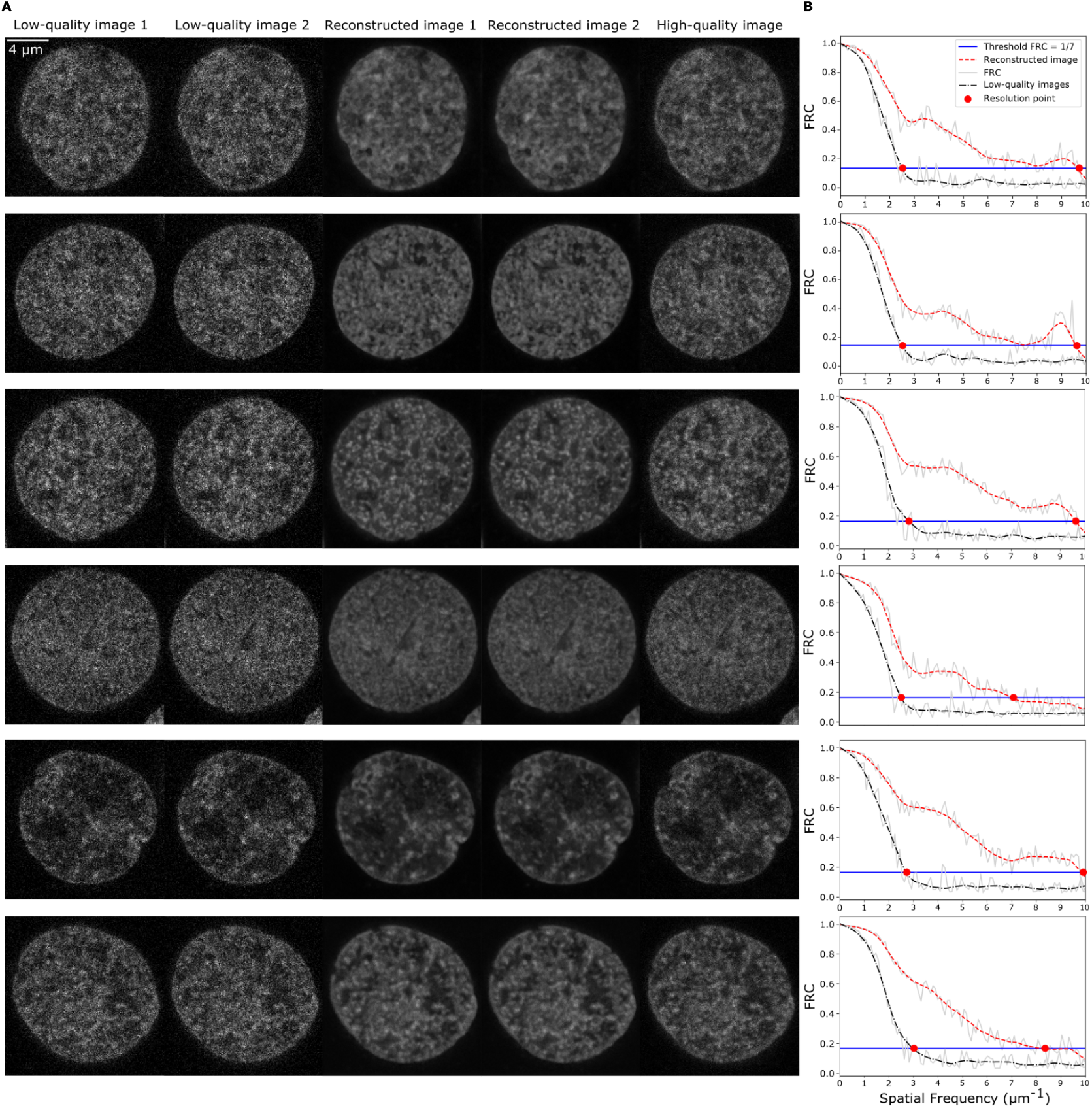
Fourier ring correlation can quantify improvements in effective image resolution obtained by Noise2Void reconstruction. A) DNA distribution in micrographs of nuclei in a fixed zebrafish embryo, obtained by stimulated emission depletion (STED) microscopy. Low quality image 1 and 2 are acquired with identical acquisition settings. Reconstructed image 1 and 2 are obtained by Noise2Void reconstruction from the low quality images 1 and 2, respectively. The high quality image was acquired in the same scanning sequence as the low quality images, but included accumulation by repeated line-scanning to improve image quality. B) Fourier ring correlation (FRC) analysis to determine the improvement in effective resolution of reconstructed images relative to the unprocessed low quality images.

**Fig S3.**
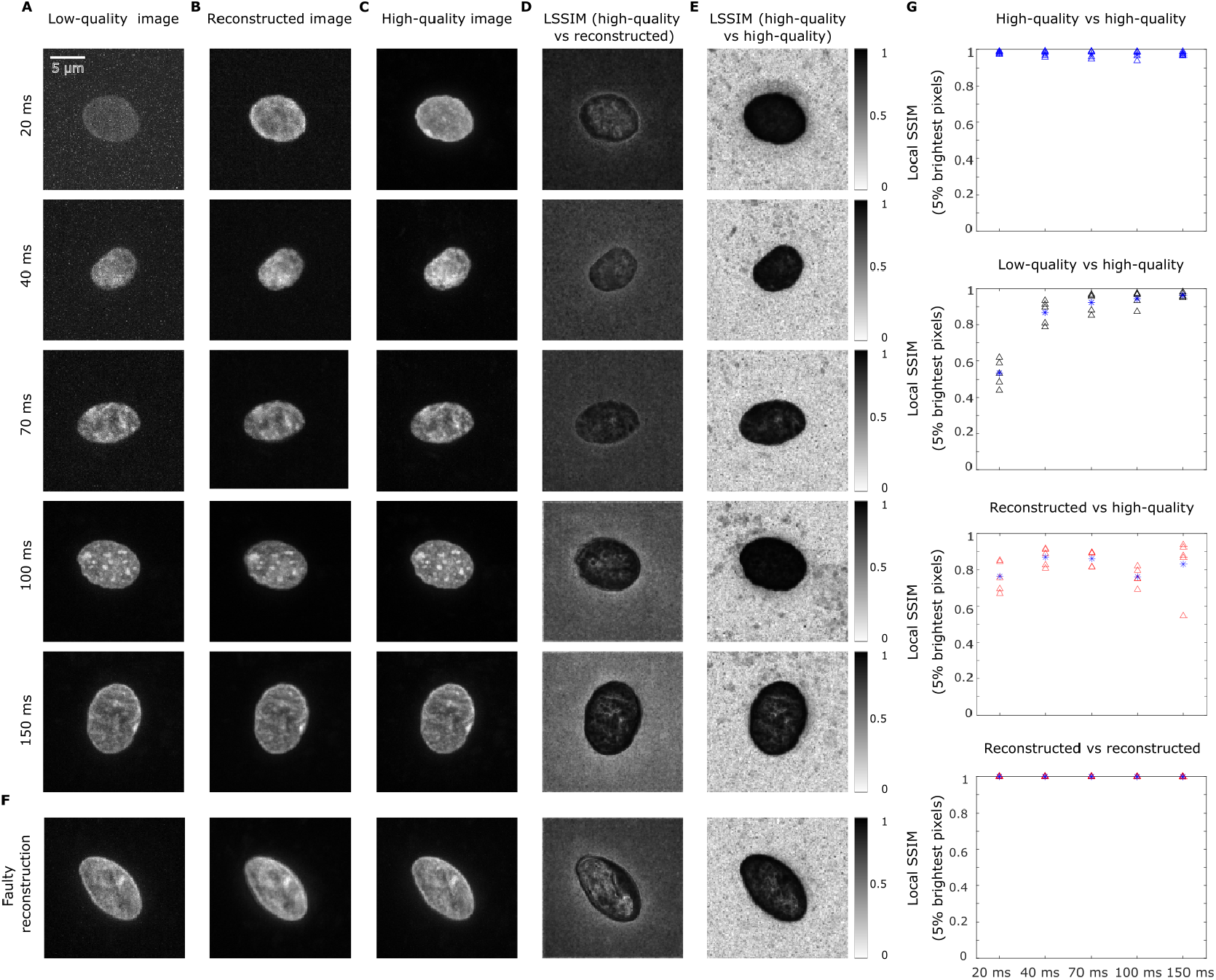
Local structural similarity index metric can show faulty reconstructions. A) Representative micrographs of nuclei of human cheek cells in which DNA was labelled by Hoechst 33342. Images are maximum intensity projections of full volumetric stacks acquired with different exposure time (*t*_*exp*_) as indicated B) Noise2Void-processed images corresponding to panel A. C) High-quality images acquired at the same position, but with *t*_*exp*_ = 200 ms. D) Local structural similarity index metric (SSIM) map for the comparison between reconstructed images and high-quality images. E) Local SSIM map for the comparison between two high-quality images acquired at the same position, suggesting that there is no structural mismatch in the area of interest. F) An example of a faulty reconstruction, indicated by a structural mismatch inside the area of interest. G) Average SSIM values based on 5% lowest local SSIM value of the 5% brightest pixels. *n* = 4, 5, 5 values from *N* = 5 different nuclei.

**Fig S4.**
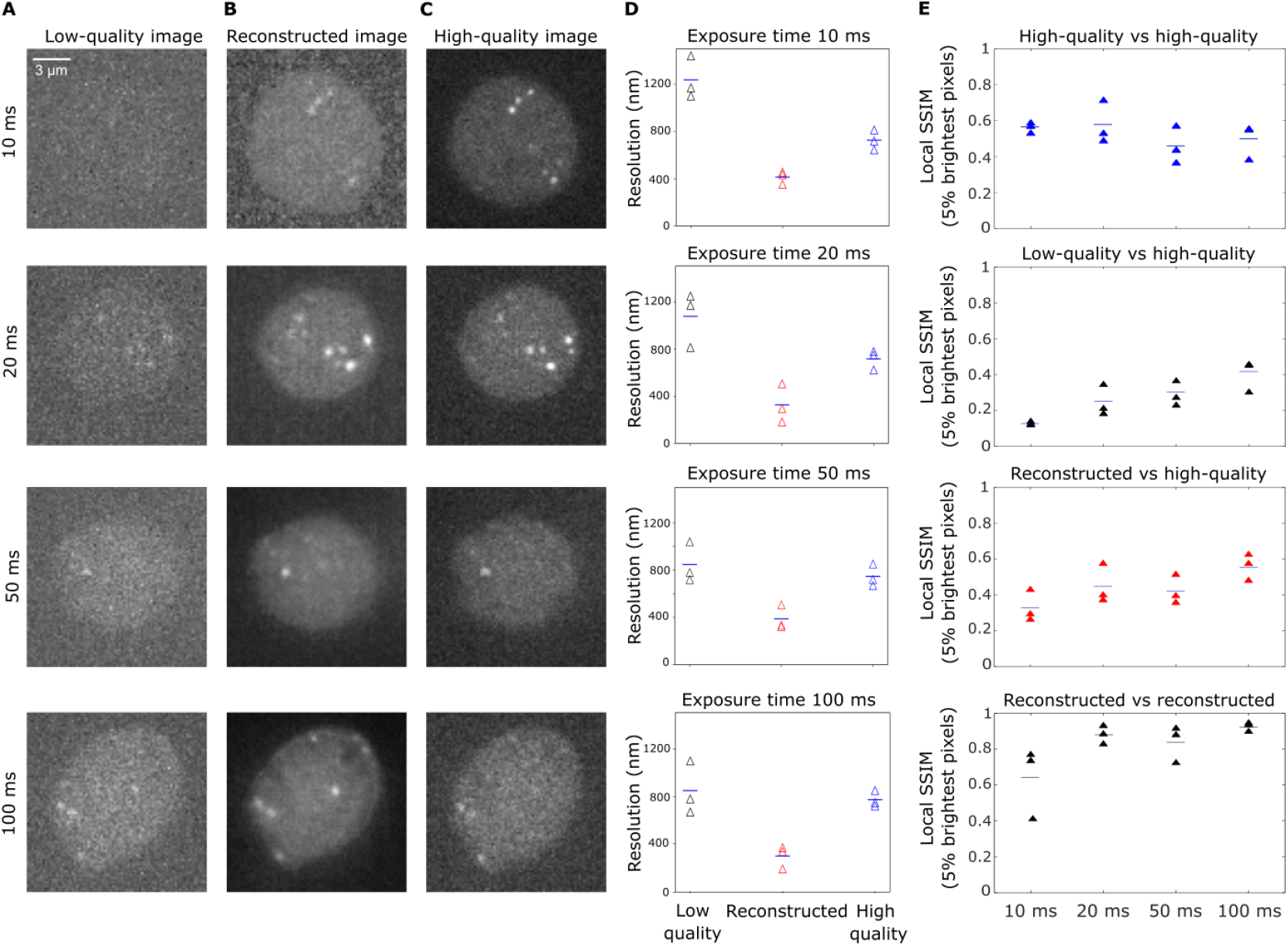
Metric-based assessment to show how far the signal-to-noise of the image can be compromised while still allowing Noise2Void recover signal-to-noise ratio post-acquisition. A) Representative micrographs of recruited RNA polymerase II (Pol II Ser5P) in live sphere-stage zebrafish embryos, visualized with antibody fragments (Fab) labelled with Janelia fluor 647. Images are single image planes and were acquired with different exposure times *t*_*exp*_ as indicated. Intensity scale from black to white adjusted to the 0.01-th and the 99.99-th percentile B) Corresponding images after Noise2Void-based reconstruction. C) Corresponding high quality reference images captured with an *t*_*exp*_ of 200 ms. D) Effective resolution as determined by FRC analysis for low quality images, reconstructed images, and high quality images for the indicated *t*_*exp*_. *n* = 4, 3, 3 values from *N* = 3 different embryos are shown with the mean. E) Average SSIM values based on the 5% lowest local SSIM values of the 5% brightest pixels. *n* = 4, 4, 3 values from *N* = 3 different embryos are shown with the mean.

**Fig S5.**
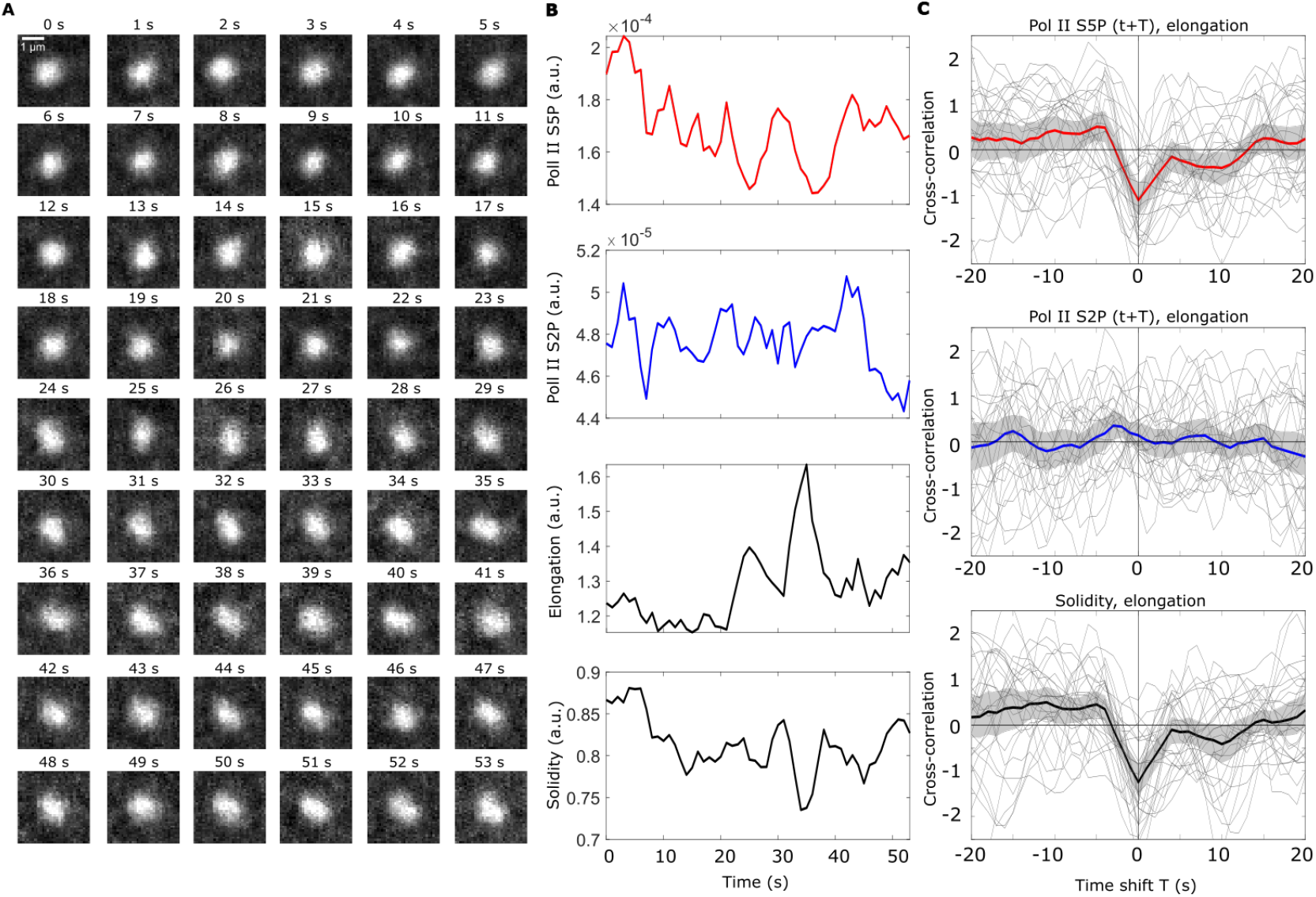
Coordinated changes in RNA polymerase cluster phosphorylation and shape are reproduced by Noise2Void-accelerated imaging with a different exposure time. A) Representative series of time-lapse images showing a single RNA polymerase II cluster in the Pol II Ser5p channel (single image plane from the middle z position of the cluster, exposure time *t*_*exp*_ = 20 ms, effective time resolution for acquisition of full 3D volumes 1 s). B) Time courses of the Pol II Ser5P intensity, the Pol II Ser2P intensity, elongation, and solidity for the example track shown in panel A. C) Cross-correlation analysis Gray lines indicate the analysis results for individual cluster time-courses, thick lines the mean over all analyzed clusters, the gray region is the bootstrap 95% confidence interval. Analysis based on *n* = 27 clusters, recorded from one sphere-stage embryo.

**Fig S6.**
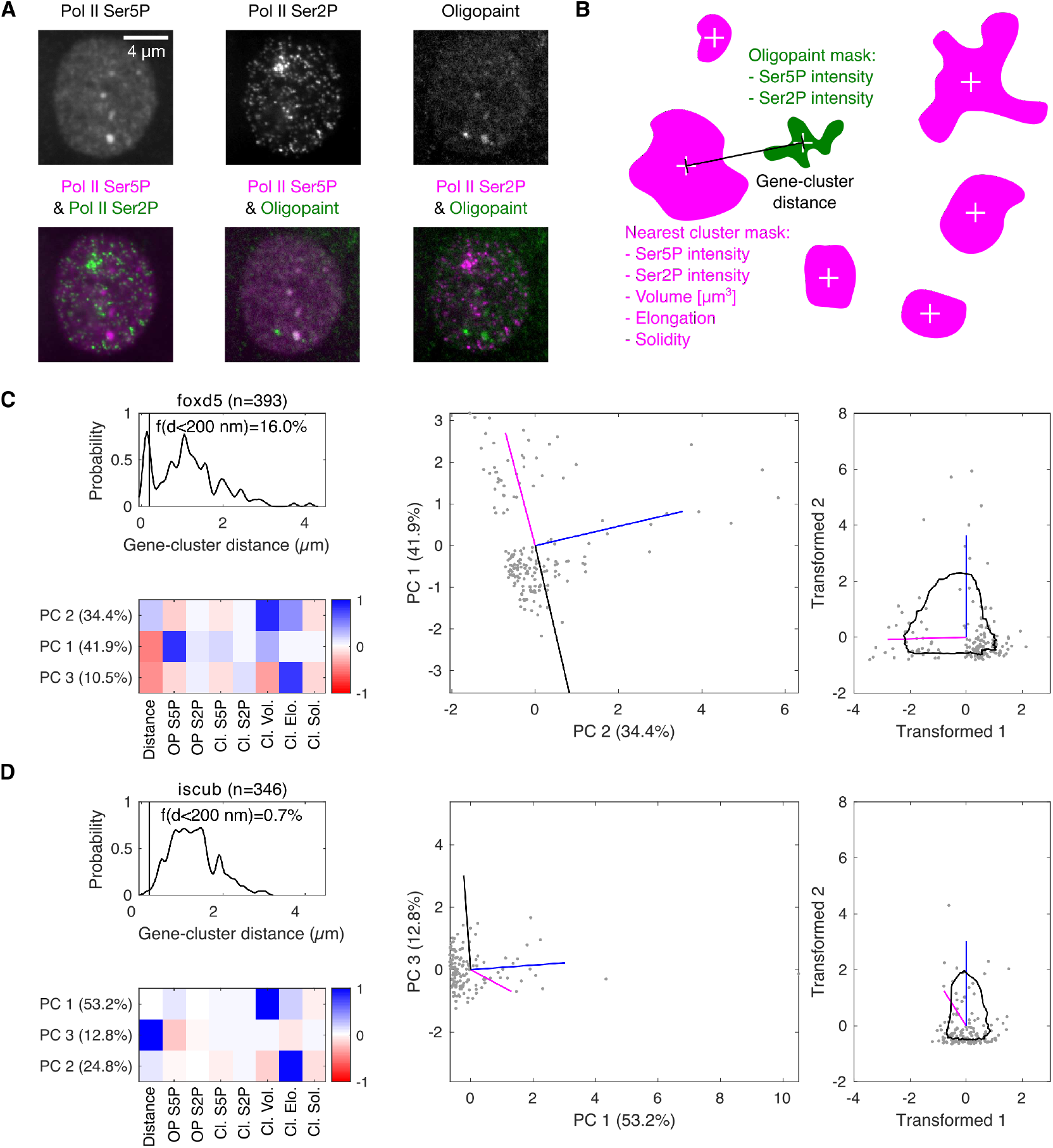
Single images of clusters from fixed embryos can be sorted in pseudo-time based on their interaction with visiting genes. A) Example micrographs of a nucleus of a fixed sphere-stage zebrafish embryo with Pol II Ser5P, Pol II Ser2P, and oligopaint fluorescence *in situ* hybridization (target gene *foxd5*) signal. B) Sketch of properties extracted from Pol II Ser5P cluster-oligopaint nearest neighbor pairs. C) Overview of the pseudp-time reconstruction procedure in the case of a gene with a high frequency of visiting Pol II Ser5P clusters (quantified as the fraction *f* of observations with less than 200 nm distance between the oligopaint signal and the nearest Pol II Ser5P cluster, *d*). The top three principal component (PC) support vectors are displayed (top two PC vectors are sorted so that the top vector has the higher weight in the volume dimension). The two top vectors are used to plot single observations in a two-dimensional overview plot. A rotation and inversion transformation are then applied to ensure the always the volume points exactly North, and the oligopaints Pol II Ser5P intensity towards the West half of the graph. Values can now be sorted according to angle relative to the direction North, and a running average over these angles indicates how well the sorted values are distributed away from the coordinate origin. In the case of the gene *foxd5*, a clear separation away from the origin can be seen, indicating a successful sorting by pseudo-time. D) In the case of the gene *iscub*, which does not frequently associate closely with Pol II Ser5P clusters, the individual points form a single cloud close to the origin and the running average line is also close to the coordinate origin, indicating that the attempt of sorting by a pseudo-time coordinate is not successful.

**Fig S7.**
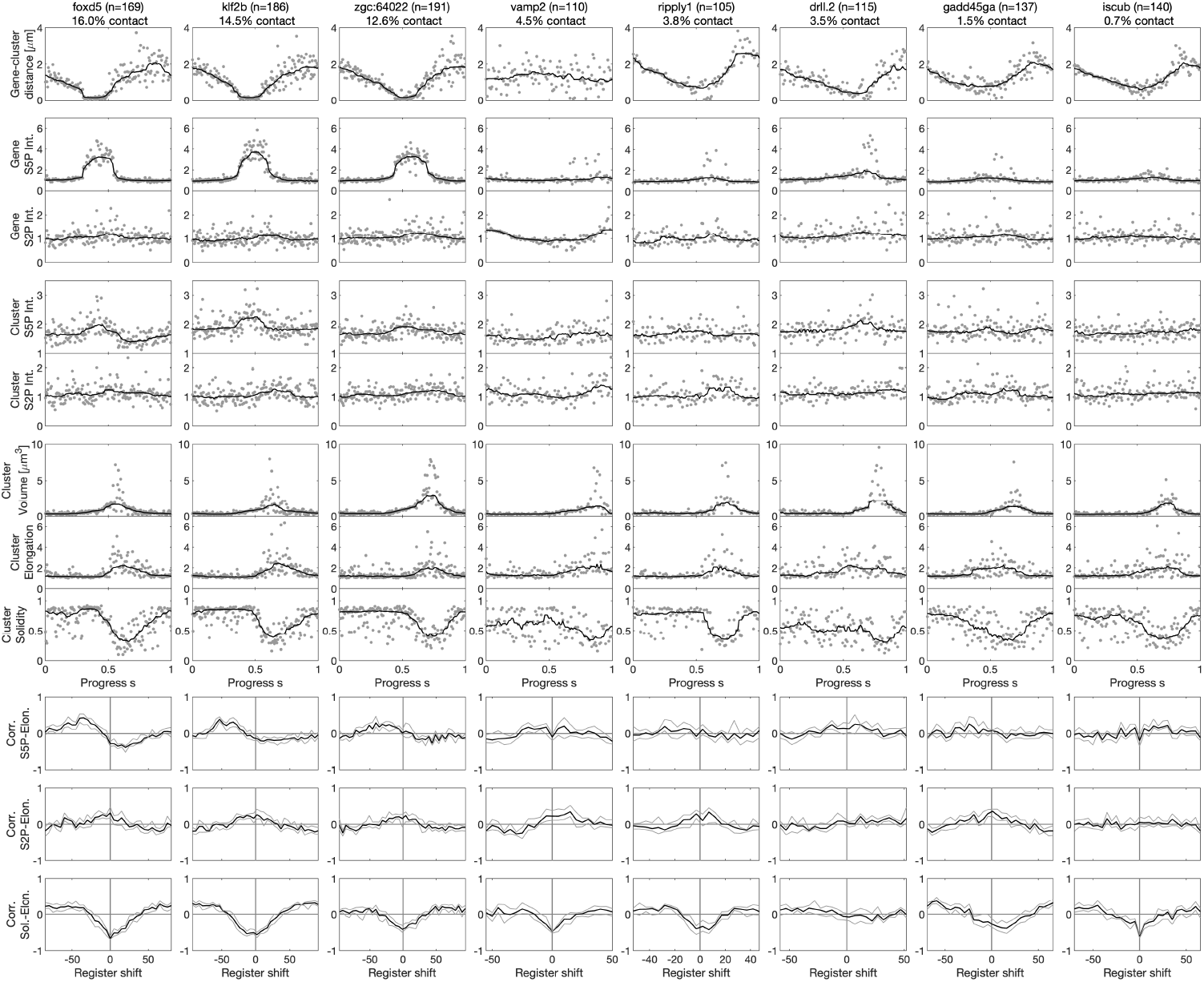
Pseudo-time sorting reproduces correlation analysis results only for genes that frequently associate with RNA polymerase II clusters. Results of the pseudo-time sorting for eight genes that were labeled oligopaint fluorescence *in-situ* hybridization. Percentage of contact (*f*) is calculated as the percentage of OP-cluster pairs with distance *d* of 200 nm or less. Shown are the pseudo-time sorted oligopaint-cluster distance, the Pol II Ser5P and Ser2P signals at the oligopaint-labeled gene (OP S5P, OP S2P, normalized against the whole nucleus median intensity), the Pol II Ser5P and Ser2P intensity at the nearest Pol II Ser5P cluster (Cl. S5P, Cl. S2P), cluster volume (Cl. Vol.), cluster elongation (Cl. Elo.), and cluster solidity (Cl. Sol.). The coordinate *s* represents the progress in pseudo-time. A register shift in pseudo-time was used to calculate the cross-correlation between cluster Pol II Ser5P intensity and elongation, cluster Pol II Ser2P intensity and elongation, and cluster solidity and elongation. Number of cluster-gene pairs included in the analysis indicated as *n* for each gene. For each gene, images were recorded from four samples, distributed over two independent experiments.

## References

1. David J. Stephens and Victoria J. Allan. Light microscopy techniques for live cell imaging. Science, 300(5616):82–86, 2003.

2. Jaroslav Icha, Michael Weber, Jennifer C. Waters, and Caren Norden. Phototoxicity in live fluorescence microscopy, and how to avoid it. BioEssays, 39(8):1700003, 2017.

3. P. Philippe Laissue, Rana A. Alghamdi, Pavel Tomancak, Emmanuel G. Reynaud, and Hari Shroff. Assessing phototoxicity in live fluorescence imaging. Nature Methods, 14(7):657–661, 2017.

4. Nicole Kilian, Alexander Goryaynov, Mark D. Lessard, Giles Hooker, Derek Toomre, James E. Rothman, and Joerg Bewersdorf. Assessing photodamage in live-cell STED microscopy. Nature Methods, 15(10):755–756, 2018.

5. James Pawley. Handbook of biological confocal microscopy, volume 236. Springer Science & Business Media, 2006.

6. Nico Scherf and Jan Huisken. The smart and gentle microscope. Nature Biotechnology, 33(8):815–818, 2015.

7. Asm Shihavuddin, Sreetama Basu, Elton Rexhepaj, Felipe Delestro, Nikita Menezes, Séverine M. Sigoillot, Elaine Del Nery, Fekrije Selimi, Nathalie Spassky, and Auguste Genovesio. Smooth 2D manifold extraction from 3D image stack. Nature Communications, 8(1):1–8, 2017.

8. William Hadley Richardson. Bayesian-based iterative method of image restoration. Journal of the Optical Society of America, 62(1):55–59, 1972.

9. Stephan Preibisch, Fernando Amat, Evangelia Stamataki, Mihail Sarov, Robert H. Singer, Eugene Myers, and Pavel Tomancak. Efficient bayesian-based multiview deconvolution. Nature Methods, 11(6):645–648, 2014.

10. Antoni Buades, Bartomeu Coll, and Jean-Michel Morel. A non-local algorithm for image denoising. In 2005 IEEE Computer Society Conference on Computer Vision and Pattern Recognition (CVPR’05), volume 2, pages 60–65. IEEE, 2005.

11. Bhawna Goyal, Ayush Dogra, Sunil Agrawal, B.S. Sohi, and Apoorav Sharma. Image denoising review: From classical to state-of-the-art approaches. Information Fusion, 55:220–244, 2020.

12. Niall O’Mahony, Sean Campbell, Anderson Carvalho, Suman Harapanahalli, Gustavo Velasco Hernandez, Lenka Krpalkova, Daniel Riordan, and Joseph Walsh. Deep learning vs. traditional computer vision. In Science and Information Conference, pages 128–144. Springer, 2019.

13. Dai Kusumoto and Shinsuke Yuasa. The application of convolutional neural network to stem cell biology. Inflammation and Regeneration, 39(1):1–7, 2019.

14. Devin P. Sullivan, Casper F. Winsnes, Lovisa Åkesson, Martin Hjelmare, Mikaela Wiking, Rutger Schutten, Linzi Campbell, Hjalti Leifsson, Scott Rhodes, Andie Nordgren, Kevin Smith, Bernard Revaz, Bergur Finnbogason, Attila Szantner, and Emma Lundberg. Deep learning is combined with massive-scale citizen science to improve large-scale image classification. Nature Biotechnology, 36(9):820–828, 2018.

15. Chawin Ounkomol, Sharmishtaa Seshamani, Mary M. Maleckar, Forrest Collman, and Gregory R. Johnson. Label-free prediction of three-dimensional fluorescence images from transmitted-light microscopy. Nature Methods, 15(11):917–920, 2018.

16. William Jones, Kaur Alasoo, Dmytro Fishman, and Leopold Parts. Computational biology: deep learning. Emerging Topics in Life Sciences, 1(3):257–274, 2017.

17. Thorsten Beier, Constantin Pape, Nasim Rahaman, Timo Prange, Stuart Berg, Davi D. Bock, Albert Cardona, Graham W. Knott, Stephen M. Plaza, Louis K. Scheffer, et al. Multicut brings automated neurite segmentation closer to human performance. Nature Methods, 14(2):101–102, 2017.

18. Tim-Oliver Buchholz, Mareike Jordan, Gaia Pigino, and Florian Jug. Cryo-care: content-aware image restoration for cryo-transmission electron microscopy data. In 2019 IEEE 16th International Symposium on Biomedical Imaging (ISBI 2019), pages 502–506. IEEE, 2019.

19. Alexander Krull, Tomáš Vičar, Mangal Prakash, Manan Lalit, and Florian Jug. Probabilistic Noise2Void: unsupervised content-aware denoising. Frontiers in Computer Science, 2:5, 2020.

20. Yair Rivenson, Zoltán Göröcs, Harun Günaydin, Yibo Zhang, Hongda Wang, and Aydogan Ozcan. Deep learning microscopy. Optica, 4(11):1437–1443, 2017.

21. Elias Nehme, Lucien E. Weiss, Tomer Michaeli, and Yoav Shechtman. Deep-STORM: super-resolution single-molecule microscopy by deep learning. Optica, 5 (4):458–464, 2018.

22. Martin Weigert, Uwe Schmidt, Tobias Boothe, Andreas Müller, Alexandr Dibrov, Akanksha Jain, Benjamin Wilhelm, Deborah Schmidt, Coleman Broaddus, Siân Culley, et al. Content-aware image restoration: pushing the limits of fluorescence microscopy. Nature Methods, 15(12):1090–1097, 2018.

23. Chinmay Belthangady and Loic A. Royer. Applications, promises, and pitfalls of deep learning for fluorescence image reconstruction. Nature Methods, 16(12): 1215–1225, 2019.

24. Erick Moen, Dylan Bannon, Takamasa Kudo, William Graf, Markus Covert, and David Van Valen. Deep learning for cellular image analysis. Nature Methods, 16 (12):1233–1246, 2019.

25. Jöel Lefebvre, Avelino Javer, Mariia Dmitrieva, Jens Rittscher, Bohdan Lewków, Edward Allgeyer, George Sirinakis, and Daniel St Johnston. Single-molecule localization microscopy reconstruction using Noise2Noise for super-resolution imaging of actin filaments. In 2020 IEEE 17th International Symposium on Biomedical Imaging (ISBI), pages 1596–1599. IEEE, 2020.

26. Jaakko Lehtinen, Jacob Munkberg, Jon Hasselgren, Samuli Laine, Tero Karras, Miika Aittala, and Timo Aila. Noise2Noise: Learning image restoration without clean data. arXiv, 2018.

27. Alexander Krull, Tim-Oliver Buchholz, and Florian Jug. Noise2Void-learning denoising from single noisy images. In Proceedings of the IEEE Conference on Computer Vision and Pattern Recognition, pages 2129–2137, 2019.

28. Zhou Wang, Alan C. Bovik, Hamid R. Sheikh, and Eero P. Simoncelli. Image quality assessment: from error visibility to structural similarity. IEEE Transactions on Image Processing, 13(4):600–612, 2004.

29. Dominique Brunet, Edward R. Vrscay, and Zhou Wang. On the mathematical properties of the structural similarity index. IEEE Transactions on Image Processing, 21(4):1488–1499, 2011.

30. Hongda Wang, Yair Rivenson, Yiyin Jin, Zhensong Wei, Ronald Gao, Harun Günaydin, Laurent A. Bentolila, Comert Kural, and Aydogan Ozcan. Deep learning enables cross-modality super-resolution in fluorescence microscopy. Nature Methods, 16(1):103–110, 2019.

31. Giorgio Tortarolo, Marco Castello, Alberto Diaspro, Sami Koho, and Giuseppe Vicidomini. Evaluating image resolution in stimulated emission depletion microscopy. Optica, 5(1):32–35, 2018.

32. Niccolo Banterle, Khanh Huy Bui, Edward A. Lemke, and Martin Beck. Fourier ring correlation as a resolution criterion for super-resolution microscopy. Journal of Structural Biology, 183(3):363–367, 2013.

33. Andrew G. York, Panagiotis Chandris, Damian Dalle Nogare, Jeffrey Head, Peter Wawrzusin, Robert S. Fischer, Ajay Chitnis, and Hari Shroff. Instant super-resolution imaging in live cells and embryos via analog image processing. Nature Methods, 10(11):1122–1126, 2013.

34. Yuko Sato, Lennart Hilbert, Haruka Oda, Yinan Wan, John M. Heddleston, Teng Leong Chew, Vasily Zaburdaev, Philipp Keller, Timothee Lionnet, Nadine L. Vastenhouw, and Hiroshi Kimura. Histone H3K27 acetylation precedes active transcription during zebrafish zygotic genome activation as revealed by live-cell analysis. Development, 146(19):dev179127, 2019.

35. Lennart Hilbert, Yuko Sato, Ksenia Kuznetsova, Tommaso Bianucci, Hiroshi Kimura, Frank Jülicher, Alf Honigmann, Vasily Zaburdaev, and Nadine L. Vastenhouw. Transcription organizes euchromatin via microphase separation. Nature Communications, 12(1):1–12, 2021.

36. Agnieszka Pancholi, Tim Klingberg, Weichun Zhang, Roshan Prizak, Irina Mamontova, Amra Noa, Marcel Sobucki, Andrei Yu Kobitski, Gerd Ulrich Nienhaus, Vasily Zaburdaev, and Lennart Hilbert. RNA polymerase II clusters form in line with surface condensation on regulatory chromatin. Molecular Systems Biology, 17(9):1–26, 2021.

37. Jonathan E. Henninger, Ozgur Oksuz, Krishna Shrinivas, Ido Sagi, Gary LeRoy, Ming M. Zheng, J. Owen Andrews, Alicia V. Zamudio, Charalampos Lazaris, Nancy M. Hannett, Tong Ihn Lee, Phillip A. Sharp, Ibrahim I. Cissé, Arup K. Chakraborty, and Richard A. Young. RNA-mediated feedback control of transcriptional condensates. Cell, 184(1):207–225.e24, 2021.

38. Linda S. Forero-Quintero, William Raymond, Tetsuya Handa, Matthew N. Saxton, Tatsuya Morisaki, Hiroshi Kimura, Edouard Bertrand, Brian Munsky, and Timothy J. Stasevich. Live-cell imaging reveals the spatiotemporal organization of endogenous RNA polymerase II phosphorylation at a single gene. Nature Communications, 12(1):1–16, 2021.

39. Markus Mund, Johannes Albertus van der Beek, Joran Deschamps, Serge Dmitrieff, Philipp Hoess, Jooske Louise Monster, Andrea Picco, François Nédélec, Marko Kaksonen, and Jonas Ries. Systematic nanoscale analysis of endocytosis links efficient vesicle formation to patterned actin nucleation. Cell, 174(4):884–896.e17, 2018.

40. Markus Mund, Aline Tschanz, Yu-Le Wu, Felix Frey, Johanna L. Mehl, Marko Kaksonen, Ori Avinoam, Ulrich S. Schwarz, and Jonas Ries. Superresolution microscopy reveals partial preassembly and subsequent bending of the clathrin coat during endocytosis. bioRxiv, 2021.

41. Joshua Batson and Loic Royer. Noise2Self: Blind denoising by self-supervision. In International Conference on Machine Learning, pages 524–533. PMLR, 2019.

42. Wesley Khademi, Sonia Rao, Clare Minnerath, Guy Hagen, and Jonathan Ventura. Self-supervised poisson-gaussian denoising. In Proceedings of the IEEE/CVF Winter Conference on Applications of Computer Vision, pages 2131–2139, 2021.

43. Samuli Laine, Tero Karras, Jaakko Lehtinen, and Timo Aila. High-quality selfsupervised deep image denoising. Advances in Neural Information Processing Systems, 32:6970–6980, 2019.

44. Anna S. Goncharova, Alf Honigmann, Florian Jug, and Alexander Krull. Improving blind spot denoising for microscopy. In Adrien Bartoli and Andrea Fusiello, editors, Computer Vision – ECCV 2020 Workshops, pages 380–393, Cham, 2020. Springer International Publishing.

45. Hamid R. Sheikh and Alan C. Bovik. Image information and visual quality. IEEE Transactions on Image Processing, 15(2):430–444, 2006.

46. Feng Wang, Trond R. Henninen, Debora Keller, and Rolf Erni. Noise2Atom: unsupervised denoising for scanning transmission electron microscopy images. Applied Microscopy, 50(1):1–9, 2020.

47. Zhou Wang and Alan C. Bovik. Reduced-and no-reference image quality assessment. IEEE Signal Processing Magazine, 28(6):29–40, 2011.

48. Lixiong Liu, Bao Liu, Hua Huang, and Alan Conrad Bovik. No-reference image quality assessment based on spatial and spectral entropies. Signal Processing: Image Communication, 29(8):856–863, 2014.

49. Siân Culley, David Albrecht, Caron Jacobs, Pedro Matos Pereira, Christophe Leterrier, Jason Mercer, and Ricardo Henriques. Quantitative mapping and minimization of super-resolution optical imaging artifacts. Nature Methods, 15(4):263–266, 2018.

50. Won-Ki Cho, Namrata Jayanth, Brian P. English, Takuma Inoue, J Owen Andrews, William Conway, Jonathan B. Grimm, Jan-Hendrik Spille, Luke D. Lavis, Timothée Lionnet, et al. RNA polymerase ii cluster dynamics predict mRNA output in living cells. eLife, 5:e13617, 2016.

51. Ibrahim I. Cisse, Ignacio Izeddin, Sebastien Z. Causse, Lydia Boudarene, Adrien Senecal, Leila Muresan, Claire Dugast-Darzacq, Bassam Hajj, Maxime Dahan, and Xavier Darzacq. Real-Time Dynamics of RNA Polymerase II Clustering in Live Human Cells. Science, 245:664–667, 2013.

52. Timothy J. Stasevich, Yoko Hayashi-Takanaka, Yuko Sato, Kazumitsu Maehara, Yasuyuki Ohkawa, Kumiko Sakata-Sogawa, Makio Tokunaga, Takahiro Nagase, Naohito Nozaki, James G. McNally, and Hiroshi Kimura. Regulation of RNA polymerase II activation by histone acetylation in single living cells. Nature, 516 (7530):272–275, 2014.

53. Barbara Steurer, Roel C. Janssens, Bart Geverts, Marit E. Geijer, Franziska Wienholz, Arjan F. Theil, Jiang Chang, Shannon Dealy, Joris Pothof, Wiggert A. Van Cappellen, Adriaan B. Houtsmuller, and Jurgen A. Marteijn. Live-cell analysis of endogenous GFP-RPB1 uncovers rapid turnover of initiating and promoter-paused RNA polymerase II. Proceedings of the National Academy of Sciences of the United States of America, 115(19):E4368–E4376, 2018.

54. Jieru Li, Ankun Dong, Kamola Saydaminova, Hill Chang, Guanshi Wang, Hiroshi Ochiai, Takashi Yamamoto, and Alexandros Pertsinidis. Single-molecule nanoscopy elucidates RNA polymerase II transcription at single genes in live cells. Cell, 178 (2):491–506.e28, 2019.

55. Jieru Li, Angela Hsu, Yujing Hua, Guanshi Wang, Lingling Cheng, Hiroshi Ochiai, Takashi Yamamoto, and Alexandros Pertsinidis. Single-gene imaging links genome topology, promoter–enhancer communication and transcription control. Nature Structural & Molecular Biology 2020, 27(11):1032–1040, 2020.

56. Yad Ghavi-Helm, Felix A. Klein, Tibor Pakozdi, Lucia Ciglar, Daan Noordermeer, Wolfgang Huber, and Eileen E. M. Furlong. Enhancer loops appear stable during development and are associated with paused polymerase. Nature, 521(7):96–100, 2014.

57. Sergio Martin Espinola, Markus Götz, Maelle Bellec, Olivier Messina, Jean Bernard Fiche, Christophe Houbron, Matthieu Dejean, Ingolf Reim, Andrés M. Cardozo Gizzi, Mounia Lagha, and Marcelo Nollmann. Cis-regulatory chromatin loops arise before TADs and gene activation, and are independent of cell fate during early Drosophila development. Nature Genetics, 53(4):477–486, 2021.

58. Eileen E.M. Furlong and Michael Levine. Developmental enhancers and chromo-some topology. Science, 361(6409):1341–1345, 2018.

59. Douglas R. Higgs. Enhancer-promoter interactions and transcription. Nature Genetics, 52(5):470–471, 2020.

60. Hugo B. Brandão, Michele Gabriele, and Anders S. Hansen. Tracking and interpreting long-range chromatin interactions with super-resolution live-cell imaging. Current Opinion in Cell Biology, 70:18–26, 2021.

61. Joshua D. Larkin, Argyris Papantonis, Peter R. Cook, and Davide Marenduzzo. Space exploration by the promoter of a long human gene during one transcription cycle. Nucleic Acids Research, 41(4):2216–27, 2013.

62. Iain Williamson, Laura A. Lettic, Robert E. Hill, and Wendy A. Bickmore. Shh and ZRS enhancer colocalisation is specific to the zone of polarising activity. Development, 143(16):2994–3001, 2016.

63. Michael I. Robson, Alessa R. Ringel, and Stefan Mundlos. Regulatory landscaping: How enhancer-promoter communication is sculpted in 3D. Molecular Cell, 74(6):1110–1122, 2019.

64. Hongtao Chen, Michal Levo, Lev Barinov, Miki Fujioka, James B. Jaynes, and Thomas Gregor. Dynamic interplay between enhancer-promoter topology and gene activity. Nature Genetics, 50(9):1296–1303, 2018.

65. Jeffrey M. Alexander, Juan Guan, Bingkun Li, Lenka Maliskova, Michael Song, Yin Shen, Bo Huang, Stavros Lomvardas, and Orion D. Weiner. Live-cell imaging reveals enhancer-dependent sox2 transcription in the absence of enhancer proximity. eLife, 8:1–42, 2019.

66. Jordan Yupeng Xiao, Antonina Hafner, and Alistair N. Boettiger. How subtle changes in 3D structure can create large changes in transcription. eLife, 10:e64320, 2021.

67. Jessica Zuin, Gregory Roth, Yinxiu Zhan, Julie Cramard, Josef Redolfi, Ewa Piskadlo, Pia Mach, Mariya Kryzhanovska, Gergely Tihanyi, Hubertus Kohler, Peter Meister, Sebastien Smallwood, and Luca Giorgetti. Nonlinear control of transcription through enhancer-promoter interactions. bioRxiv, 2021.

68. Christopher H. Eskiw, Alexander Rapp, David R. F. Carter, and Peter R. Cook. RNA polymerase II activity is located on the surface of protein-rich transcription factories. Journal of Cell Science, 121(12):1999–2007, 2008.

69. Ryu-Suke Nozawa, Lora Boteva, Dinesh C. Soares, Catherine Naughton, Alison R. Dun, Adam Buckle, Bernard Ramsahoye, Peter C. Bruton, Rebecca S. Saleeb, Maria Arnedo, Bill Hill, Rory R. Duncan, Sutherland K. Maciver, and Nick Gilbert. SAF-A regulates interphase chromosome structure through oligomerization with chromatin-associated RNAs. Cell, 169:1214–1227, 2017.

70. Yefei Yin, Yuyang J. Lu, Xuechun Zhang, Wen Shao, Yanhui Xu, Pan Li, Yantao Hong, Li Cui, Ge Shan, Bin Tian, Qiangfeng Cliff Zhang, and Xiaohua Shen. U1 snRNP regulates chromatin retention of noncoding RNAs. Nature, 580:147–150, 2020.

71. Wen Shao, Xianju Bi, Boyang Gao, Jun Wu, Yixuan Pan, Yafei Yin, Zhimin Liu, Wenhao Zhang, Xu Jiang, Wenlin Ren, Yanhui Xu, Zhongyang Wu, Kaili Wang, Ge Zhan, J. Yuyang Lu, Xue Han, Tong Li, Jianlong Wang, Guohong Li, Haiteng Deng, Bing Li, and Xiaohua Shen. Phase separation of RNA-binding protein promotes polymerase engagement and transcription. bioRxiv, page 2021.03.26.436939, 2021.

72. Tomas Pachano, Víctor Sánchez-Gaya, Thais Ealo, Maria Mariner-Faulí, Tore Bleckwehl, Helena G. Asenjo, Patricia Respuela, Sara Cruz-Molina, María Muñoz-San Martín, Endika Haro, Wilfred F.J. van IJcken, David Landeira, and Alvaro Rada-Iglesias. Orphan CpG islands amplify poised enhancer regulatory activity and determine target gene responsiveness. Nature Genetics, 53(7):1036–1049, 2021.

73. Amra Noa, Hui Shun Kuan, Vera Aschmann, Vasily Zaburdaev, and Lennart Hilbert. The hierarchical packing of euchromatin domains can be described as multiplicative cascades. PLoS Computational Biology, 17(5):1–16, 2021.

74. Ye Zhang, Kenian Chen, Steven A. Sloan, Mariko L. Bennett, Anja R. Scholze, Sean O’Keeffe, Hemali P. Phatnani, Paolo Guarnieri, Christine Caneda, Nadine Ruderisch, et al. An RNA-sequencing transcriptome and splicing database of glia, neurons, and vascular cells of the cerebral cortex. Journal of Neuroscience, 34 (36):11929–11947, 2014.

75. John M. Gaspar. Improved peak-calling with MACS2. BioRxiv, 2018.

76. Elizabeth J.F. White, Aerielle E. Matsangos, and Gerald M. Wilson. AUF1 regulation of coding and noncoding RNA. Wiley Interdisciplinary Reviews: RNA, 8(2):e1393, 2017.

77. Guy Nir, Irene Farabella, Cynthia Pérez Estrada, Carl G. Ebeling, Brian J. Beliveau, Hiroshi M. Sasaki, S. Dean Lee, Son C. Nguyen, Ruth B. McCole, Shyamtanu Chattoraj, et al. Walking along chromosomes with super-resolution imaging, contact maps, and integrative modeling. PLoS Genetics, 14(12):e1007872, 2018.

