## Supplementary material for "Deep-learning microscopy image reconstruction with quality control reveals second-scale rearrangements in RNA polymerase II clusters": Scripts used for the selection of genes: oligopaints_script.m

```
clc
clear

% This script appends streets and makes Oligopaint probes to be ordered,
% given (a) which street should go to which locus for the Locus-Specific
% part, (b) Universal MS and BS, and (c) Universal Flourophore Part.

% This is useful if we already have a list of streets that work well
% together and have a partially overlapping set of genes (ROIs) that we
% previously ordered probes for.

% DESIGN %%%%%%%%%%%%%%%%%%
% UniversalForward-LocusSpecific-UniversalFlourophore-HybridizingProbe-UniversalBackward
%%%%%%%%%%%%%%%%%%%%%%%%%%%

MainStreetsIntersectedFile = strcat('oligopaints/Main_isected.txt');
GivenStreetsFilePath = strcat('oligopaints/GivenStreets.txt');
OligosSavePath = strcat('oligopaints/');
UniversalFilePath = strcat('oligopaints/Universal.txt');

% Read in the given Streets for our ROIs.
fid = fopen(GivenStreetsFilePath);
GivenStreets = textscan(fid,'%s%s%s');
fclose(fid);
% Contains Headers!
% Cell 1: Names of the ROIs
% Cell 2: Number of the LS Street in the Streets List Ref File
% Cell 3: Street sequence (RC of this should be appended!!)

% Read in the Intersected Probes for our ROIs.
fid = fopen(MainStreetsIntersectedFile);
HybridizingProbes = textscan(fid,'%s%s%s%s%s%s%s%s%s');
fclose(fid);
% Doesn't contain Headers!
% Cell 1: Chromosome of ROI
% Cell 2: Start of ROI
% Cell 3: End of ROI
% Cell 4: Name of ROI
% Cell 5: Chromosome of Probe
% Cell 6: Start of Probe
% Cell 7: End of Probe
% Cell 8: Sequence of Probe
% Cell 7: Temperature of Probe

% Read in the given Universals for our ROIs.
fid = fopen(UniversalFilePath);
tline = fgetl(fid);
tline = fgetl(fid);
Universals = strsplit(tline);
fclose(fid);
% Contains Headers!
% Cell 1: Number of the Universal Forward in the Streets List Ref File
% Cell 2: Universal Forward Sequence 
% Cell 3: Number of Original Universal Forward For Flourophore in the Ref File
% Cell 4: Original Universal Forward Sequence
% Cell 5: Number of the Universal Backward in the Streets List Ref File
% Cell 6: Universal Backward Sequence (RC of this should be appended!!) ~ Same as Primer to be ordered

OligopaintsFileID = fopen(strcat(OligosSavePath, 'Oligopaints.txt'),'w');
OligopaintSequences=strings([numel(HybridizingProbes{1}),1]);
for i=1:numel(HybridizingProbes{1})
    ROI = HybridizingProbes{4}{i};
    OligopaintName =  strcat(ROI,'_',num2str(i));
    
    for j=1:numel(GivenStreets{1})
        ROI_to_match = GivenStreets{1}{j};
        if strcmp(ROI, ROI_to_match)
            % found matching ROI
            LocusSpecificBit = ReverseComplement(GivenStreets{3}{j},1);
            break
        end
    end
    
    OligopaintSequences(i) = strcat(Universals{2}, LocusSpecificBit, Universals{4}, HybridizingProbes{8}{i}, ReverseComplement(Universals{6},1));
    fprintf(OligopaintsFileID,'%s,%s\n',OligopaintName,OligopaintSequences(i));
end
fclose(OligopaintsFileID);

ROI_names=string(HybridizingProbes{4});
ROI_names_stable=unique(ROI_names,'stable');
NumProbes_list=cell2mat(cellfun(@(x) sum(ismember(ROI_names,x)),ROI_names_stable,'un',0));
MSIDsFileID = fopen(strcat(OligosSavePath, 'MS_IDs.txt'),'w');
MSDensityFileID = fopen(strcat(OligosSavePath, 'MSDensity.txt'),'w');
fprintf(MSIDsFileID,'%s\t%s\t%s\t%s\t%s\t%s\n','MSRegionID','ID','StreetNum','StreetSequence','BridgeSequence','ForwardPrimerSequence');
fprintf(MSDensityFileID,'%s\t%s\t%s\t%s\t%s\t%s\t%s\n','MS_Region','Chromosome','Start','End','Size_(kb)','#_of_OPs_probes','Density_(Probes/kb)');
for i=2:numel(GivenStreets{1})
    fprintf(MSIDsFileID,'%s\t%s\t%s\t%s\t%s\t%s\n',GivenStreets{1}{i},num2str(i-1),GivenStreets{2}{i},GivenStreets{3}{i},GivenStreets{3}{i},ReverseComplement(GivenStreets{3}{i},1));
    
    for j=1:numel(HybridizingProbes{1})
        if strcmp(GivenStreets{1}{i}, HybridizingProbes{4}{j})
            Chromosome=HybridizingProbes{1}{j};
            Start=HybridizingProbes{2}{j};
            End=HybridizingProbes{3}{j};
            break
        end
    end
    Size=(str2double(End)-str2double(Start))/1000;
    NumProbes=NumProbes_list(i-1);
    Density=NumProbes/Size;
    fprintf(MSDensityFileID,'%s\t%s\t%s\t%s\t%s\t%s\t%s\n',GivenStreets{1}{i},Chromosome,Start,End,num2str(round(Size,0)),num2str(NumProbes),num2str(Density));
end
fclose(MSDensityFileID);
fclose(MSIDsFileID);

fileID=fopen(strcat(OligosSavePath,'PrimersToOrder.txt'),'w');
fprintf(fileID,'Locus-specific primers:\n');
fprintf(fileID,'-----------------------\n\n');
fprintf(fileID,'LocusName\tStreetNum\tForwardPrimerSeq\n');
for i=2:numel(GivenStreets{1})
    fprintf(fileID,'%s\t%s\t%s\n',GivenStreets{1}{i},GivenStreets{2}{i},ReverseComplement(GivenStreets{3}{i},1));
end 
fprintf(fileID,'\n\nUniversal primers:\n');
fprintf(fileID,'----------------------\n\n');
fprintf(fileID,'%s\t%s\t%s\n', 'UniversalForwardPrimer',Universals{1},Universals{2});
fprintf(fileID,'%s\t%s\t%s\n', 'FlourophoreContainingForwardPrimer',Universals{3},Universals{4});
fprintf(fileID,'%s\t%s\t%s\n', 'UniversalReversePrimer',Universals{5},Universals{6});
fclose(fileID);
```
