## Supplementary material for "Deep-learning microscopy image reconstruction with quality control reveals second-scale rearrangements in RNA polymerase II clusters": Scripts used for the selection of genes: RNASeq_analysis.r

```
library(rtracklayer)
library(AnnotationHub)
library(Gviz)
library(TxDb.Drerio.UCSC.danRer10.refGene)
library(org.Dr.eg.db)
library(DESeq2)
library(magrittr)
library(gplots)

# Load in the databases
dr <- org.Dr.eg.db
txdb <- TxDb.Drerio.UCSC.danRer10.refGene

# Load in the ChIP-seq data
ChIPSeq_bigWig_filepath <- "chipseq/coverage_data/Ser5P_danRer10.bigWig"
ChIPSeq_peaks_filepath <- "chipseq/peaks_data/Ser5P_danRer10_peaks.narrowPeak"
coverage = import(bigWig_filepath, format="BigWig")
peaks = import(MACS2_peaks_filepath)

# Load in RNA-seq data
RNASeq_counts_filepath <- "rnaseq/raw_counts.tsv" # downloaded from https://doi.org/10.7554/eLife.30860.025
RNASeq_sampleinfo_filepath <- "rnaseq/sample_information.tsv" # downloaded from https://doi.org/10.7554/eLife.30860.024
RNASeq_TPM_filepath <- "rnaseq/TPM.tsv" # downloaded from https://doi.org/10.7554/eLife.30860.026
counts = read.delim(RNASeq_counts_filepath, sep = "\t", dec = ".", header=T)
sample_information = read.delim(RNASeq_sampleinfo_filepath, sep = "\t", dec = ".", header=T)
TPM = read.delim(RNASeq_TPM_filepath, sep = "\t", dec = ".", header=T)

orig_genes <- genes(txdb)
dstream <- 200
ustream <- 2000
genes <- shift(orig_genes, dstream) # shifting genes so that promoter and genes don't have an overlap
promoters <- promoters(orig_genes, upstream=ustream, downstream=dstream)
entrez.ids <- genes$gene_id
promoters$SYMBOL <- mapIds(org.Dr.eg.db, keys=entrez.ids, column="SYMBOL", keytype="ENTREZID", multiVals="first")
genes$SYMBOL <- mapIds(org.Dr.eg.db, keys=entrez.ids, column="SYMBOL", keytype="ENTREZID", multiVals="first")

# filtering of peaks to include those with adjusted q-value<0.0001 and signal_value>=3
peaks_filtered <- peaks[peaks$signalValue>=3 & peaks$qValue>=4,]
mcols(peaks_filtered)$promoter <- NA
mcols(peaks_filtered)$gene <- NA
ov <- findOverlaps(peaks_filtered, promoters)
mcols(peaks_filtered)[queryHits(ov), "promoter"] <- mcols(promoters)[subjectHits(ov), "SYMBOL"]
ov <- findOverlaps(peaks_filtered, genes)
mcols(peaks_filtered)[queryHits(ov), "gene"] <- mcols(genes)[subjectHits(ov), "SYMBOL"]

promoters_with_peak <- subsetByOverlaps(promoters, peaks_filtered)

# this is a list of genes which have a Ser5P signal in their promoter (they can have a signal in their gene body too)
genes_Ser5P <- as.character(promoters_with_peak$SYMBOL)

countdata <- counts[,9:ncol(counts)]
rownames(countdata) <- counts$Gene.ID
condition <- sample_information[1:90,]
condition$stageName <- factor(condition$stageName)
condition$stageName %<>% relevel("Dome")
dds <- DESeqDataSetFromMatrix(countData = countdata,
                              colData = condition,
                              design = ~ stageName)
dds <- estimateSizeFactors(dds)
sizeFactors(dds)
normalized_counts <- counts(dds, normalized=TRUE)
write.table(normalized_counts, file="rnaseq/normalized_counts.txt", sep="\t", quote=F, col.names=NA)
ens.str <- rownames(normalized_counts)
normalized_counts <- as.data.frame(normalized_counts)
normalized_counts$Name <- mapIds(org.Dr.eg.db,
                                 keys=ens.str,
                                 column="SYMBOL",
                                 keytype="ENSEMBL",
                                 multiVals="first")

normalized_counts_Ser5P <- normalized_counts[normalized_counts$Name %in% genes_Ser5P,]
colnames(normalized_counts_Ser5P)[1:90] <- sample_information$stageName[1:90]

expression_threshold <- 0.01
fc_threshold <- 0.7 # fold change allowed between Dome and 50% epiboly
X <- Ser5P_promoter_peaks[order(-Ser5P_promoter_peaks$signalValue),]

IA <- c()
AI <- c()
II <- c()
AA <- c()

for(i in 1:length(X)){
        I = which(normalized_counts_Ser5P$Name==X$promoter[i])
        if(length(I)==0){
                next
        }
        if(length(I)>1){
                I <- I[1]
        }
        mean_expression_before_Dome <- mean(as.numeric(normalized_counts_Ser5P[I,1:20]))
        
        if(mean(as.numeric(normalized_counts_Ser5P[I,21:25]))<=expression_threshold & mean(as.numeric(normalized_counts_Ser5P[I,26:30]))<=expression_threshold){
                pdf(paste("II/", i, "_", X$promoter[i],".pdf"))
                II <- c(II, normalized_counts_Ser5P$Name[I])
        }
        if(mean(as.numeric(normalized_counts_Ser5P[I,21:25]))<=expression_threshold & mean(as.numeric(normalized_counts_Ser5P[I,26:30]))>expression_threshold){
                pdf(paste("IA/", i, "_", X$promoter[i],".pdf"))  
                IA <- c(IA, normalized_counts_Ser5P$Name[I])
        }
        if(mean(as.numeric(normalized_counts_Ser5P[I,21:25]))>expression_threshold & mean(as.numeric(normalized_counts_Ser5P[I,26:30]))>expression_threshold){
                pdf(paste("AA/", i, "_", X$promoter[i],".pdf"))    
                AA <- c(AA, normalized_counts_Ser5P$Name[I])
        }
        if(mean(as.numeric(normalized_counts_Ser5P[I,21:25]))>=expression_threshold & mean(as.numeric(normalized_counts_Ser5P[I,26:30]))<=fc_threshold*mean(as.numeric(normalized_counts_Ser5P[I,21:25]))){
                pdf(paste("AI/", i, "_", X$promoter[i],".pdf"))   
                AI <- c(AI, normalized_counts_Ser5P$Name[I])
        }
        
        plot(as.numeric(normalized_counts_Ser5P[I,1:ncol(normalized_counts_Ser5P)-1]), type="o", ylab="normalized counts")
        mylims <- par("usr")
        lines(c(1,1), mylims[3:4], col="darkgray")
        lines(c(5,5), mylims[3:4], col="darkgray")
        text(3, y = 0.9*mylims[4], labels="1-cell", srt=90, col="darkgray")
        lines(c(6,6), mylims[3:4], col="darkgray")
        lines(c(10,10), mylims[3:4], col="darkgray")
        text(8, y = 0.9*mylims[4], labels="2-cell", srt=90, col="darkgray")
        lines(c(11,11), mylims[3:4], col="darkgray")
        lines(c(15,15), mylims[3:4], col="darkgray")
        text(13, y = 0.9*mylims[4], labels="128-cell", srt=90, col="darkgray")
        lines(c(16,16), mylims[3:4], col="darkgray")
        lines(c(20,20), mylims[3:4], col="darkgray")
        text(18, y = 0.9*mylims[4], labels="1k-cell", srt=90, col="darkgray")
        lines(c(26,26), mylims[3:4], col="darkgray")
        lines(c(30,30), mylims[3:4], col="darkgray")
        text(28, y = 0.9*mylims[4], labels="50%-epiboly", srt=90, col="darkgray")
        
        lines(c(21,21), mylims[3:4], col="red")
        lines(c(25,25), mylims[3:4], col="red")
        text(23, y = 0.9*mylims[4], labels="Dome", srt=90, col="red")
        
        lines(c(1,90), c(expression_threshold, expression_threshold), col="red", lty=2)
        title(paste(normalized_counts_Ser5P$Name[I], "; peak strength=", round(X$signalValue[i],1), ", q-val=", round(X$qValue[i],1)))
        
        fc_toDome <- mean(as.numeric(normalized_counts_Ser5P[I,21:25]))/mean(as.numeric(normalized_counts_Ser5P[I,16:20]))
        fc_fromDome <- mean(as.numeric(normalized_counts_Ser5P[I,26:30]))/mean(as.numeric(normalized_counts_Ser5P[I,21:25]))
        
        mtext("fold-change", side=3, line=0, at=5)
        mtext(paste(round(fc_toDome,1)), side=3, line=0, at=18)
        mtext(paste(round(fc_fromDome,1)), side=3, line=0, at=28)
        dev.off() 
}

write.table(data.frame(II), 'II.txt', row.names = FALSE, col.names = FALSE, quote=FALSE)
write.table(data.frame(AI), 'AI.txt', row.names = FALSE, col.names = FALSE, quote=FALSE)
write.table(data.frame(AA), 'AA.txt', row.names = FALSE, col.names = FALSE, quote=FALSE)
write.table(data.frame(IA), 'IA.txt', row.names = FALSE, col.names = FALSE, quote=FALSE)
```
